# Chronic but not acute morphine exposure reversibly impairs spike generation and repetitive firing in a functionally distinct subpopulation of orexin neurons

**DOI:** 10.1101/2025.03.20.644444

**Authors:** Elizabeth A. Berry, Ellen N. Huhulea, Masaru Ishibashi, Ronald McGregor, Jerome M. Siegel, Christopher S. Leonard

**Author notes:** Corresponding Author: Christopher S. Leonard Department of Physiology New York Medical College Valhalla, NY 10595 USA.

## Abstract

Orexin (hypocretin) neuropeptides regulate numerous essential functions including sleep/wake state stability and reward processing. Orexin synthesizing neurons respond to drug cues and undergo structural changes following persistent drug exposure. Post-mortem brains from opioid users, and opioid-treated rodents have orexin somata that become ∼20 % smaller and ∼50% more numerous and are postulated to promote hyper-motivation for drug-seeking though increased orexin release. Biophysical considerations suggest that decreased soma size should increase cellular excitability, however the impact of chronic opioids on firing ability, which drives peptide release, has not been explored. To test this, we assessed the intrinsic electrophysiological properties of orexin neurons by whole-cell recordings in slices from male orexin-EGFP mice treated by daily morphine or saline injections for two weeks. Paradoxically, we found that while daily morphine decreased average soma size, it impaired excitability in a subpopulation of orexin neurons identified by electrophysiological criteria as “H-type”, while entirely sparing “D-type” neurons. This impairment was manifest by smaller, broader action potentials, variable firing and a downscaling of firing gain. These adaptations required more than a single morphine dose and recovered, along with soma size, after four weeks of passive withdrawal. Taken together, these observations indicate that daily opioid exposure differentially impacts H-type orexin neurons and predicts that the ability of these neurons to encode synaptic inputs into spike trains and to release neuropeptides becomes impaired in conjunction with opioid dependence.

**Significance Statement:** Orexin peptide signaling sustains motivation for opioid and cocaine seeking and chronic use upregulates orexin peptides and induces structural changes consistent with enhanced orexin release. However, the impact of chronic drug exposure on the ability of orexin neurons to fire action potentials which drive peptide release has not been explored. Paradoxically, we found that two weeks of daily morphine exposure selectively and reversibly impaired action potential firing in a distinct subpopulation of orexin neurons (H-cells). These findings further support a functional dichotomy among orexin neurons and imply that the ability of H-cells to encode input signals and release neuropeptides becomes impaired during development of opioid dependence.

## Introduction

Orexin (hypocretin) neuropeptides are synthesized by neurons located around the fornix in the hypothalamus (de Lecea et al., 1998; Sakurai et al., 1998) that have widespread central projections and regulate numerous functions including arousal, feeding and reward (Peyron et al., 1998; Sakurai, 2007; Mahler et al., 2014; Li, Giardino and de Lecea, 2017). Orexin peptides act through two G protein-coupled receptors, orexin 1 (OX1R) and 2 (OX2R) to depolarize and elevate intracellular calcium levels in cells expressing these receptors (Muraki et al., 2004; Kukkonen and Leonard, 2014; Leonard and Kukkonen, 2014).

Intact orexin signaling is critical for the normal regulation of sleep and waking states since genetic disruption results in narcolepsy with cataplexy (Chemelli et al., 1999; Lin et al., 1999; Willie et al., 2003; Kalogiannis et al., 2011) and patients with this sleep disorder have little, to no orexin peptide (Peyron et al., 2000; Thannickal et al., 2000; Scammell, 2003; Seifinejad et al., 2023).

Strong evidence also links the orexin system to motivation and reward processing. Many orexin neurons are located in the lateral hypothalamic area (LHA), a region classically linked to feeding, drinking and reinforcement (Stuber and Wise, 2016). Moreover, ∼50 % of orexin neurons are immunopositive for mu-opioid receptors and respond to chronic morphine exposure with increased activity of the cAMP response element (cAMPRE) and to naloxone-precipitated withdrawal with increased cAMPRE and cFOS (Georgescu et al., 2003). In brain slice experiments, about half of mouse orexin neurons are directly inhibited and show decreased EPSC frequency following acute morphine application (Li and van den Pol, 2008) and OX1R signaling in the VTA is necessary for induction of excitatory synaptic plasticity in dopamine neurons following single cocaine or morphine injections (Borgland et al., 2006; Baimel and Borgland, 2015).

Orexin neurons are also cfos-responsive to drug-related cues (Harris, Wimmer and Aston-Jones, 2005; Harris and Aston-Jones, 2006) and orexin-A can reinstate extinguished drug seeking (Boutrel et al., 2005; Harris, Wimmer and Aston-Jones, 2005), while the genetic absence of orexin dampens reward-related behavior (Georgescu et al., 2003; McGregor et al., 2011; Shaw et al., 2017) and orexin receptor antagonists attenuate drug-induced behavioral changes in rodent substance use disorder models (Harris, Wimmer and Aston-Jones, 2005; Hutcheson et al., 2011; Brown, Khoo and Lawrence, 2013; Gentile et al., 2018; James and Aston-Jones, 2020).

Consistent with participating in neural adaptations underlying addiction, orexin neurons undergo structural and synaptic plasticity in response to both environmental challenges and persistent exposure to drugs of abuse. Orexin neurons show increased numbers of asymmetric somatic synapses and increase mEPSP frequency after overnight fasting (Horvath and Gao, 2005) but show decreased synaptic integration following diet-induced obesity (Tan et al., 2020). Sleep deprivation decreases perisomatic astrocytic processes resulting in presynaptic inhibition of glutamatergic inputs (Briggs, Hirasawa and Semba, 2018), while apparently increasing the AMPA/NMDA ratio (Rao et al., 2007) – a change also observed following three days of cocaine injections which facilitated high frequency LTP (Rao et al., 2013).

Strikingly, brains from human heroin addicts and mice chronically treated with morphine have ∼50% more immunodetectable orexin neurons with ∼20% decrease in soma size (Thannickal et al., 2018) and an increased orexin fiber density and TH levels in mouse locus coeruleus (McGregor et al., 2022) and VTA (McGregor et al., 2024). Comparable increases in orexin immunopositive neurons were observed in rats following self-administration of cocaine (James et al., 2019) and fentanyl (Fragale, James and Aston-Jones, 2021) suggesting dysregulation of an “orexin reserve” that might enhance orexin release and drive hyper-motivation for drug taking (James and Aston-Jones, 2022).

However, orexin output depends not only on peptide availability, but also on orexin neuron excitability, and since the effects of chronic morphine on orexin neuron excitability were unknown, we used brain slices and whole-cell recordings to investigate the intrinsic electrical properties of orexin neurons following two weeks of daily morphine or saline injections.

## Materials and Methods

All procedures complied with NIH guidelines and were approved by New York Medical College Institutional Animal Care and Use Committee and all efforts were directed at minimizing the number of animals used for this study.

### Mice

Male orexin-EGFP mice, which express enhanced green fluorescence protein (EGFP) under the control of the human prepro-orexin promoter (Yamanaka et al., 2003), were used for all experiments. These mice were bred in our animal facility from founder mice generously provided by Dr. Luis De Lecea and Dr. William J Giardino (Adamantidis et al., 2007). For the duration of all experiments, mice were singly housed in standard mouse cages (10.5 in x 6 in x 5.75 in; 22 ± 1°C) and kept on a 12-12 light-dark cycle with lights on at five am. Mice had ad lib. access to water and food (Purina rodent laboratory diet 5001, protein 23 %, fat 4.5 %, fiber 6 %).

### Addiction, Withdrawal, 1-Day Injection paradigms

Experiments used three treatment cohorts that we termed the “Addiction”, “Withdrawal”, and “1-Day cohorts”. Mice in each cohort were randomly assigned to one of two groups that received single, daily subcutaneous injections of either sterile saline (0.9 % NaCl) or 50 mg/kg morphine sulfate dissolved in sterile saline between 9:30 – 10:30am. Mice were weighed daily, just before injection, to determine the precise volume (∼ 0.1 ml) of morphine/saline to be administered. Mice were then sacrificed for brain slice electrophysiology or for immunohistochemistry according to the schedule illustrated in Fig. 1A. Mice in the Addiction cohort received daily saline or morphine injections for 14 days and were sacrificed between 11:30am -12:30pm on the day of their last injection. Mice in the Withdrawal cohort received injections as in the Addiction paradigm but then received no injections for four-weeks. During this four-week period, mice were checked and weighed a few times per week to monitor for overall health and weight change and received weekly cage changes by veterinary staff. On the last day of the four-week period, mice were sacrificed for slice electrophysiology or immunohistochemistry, at the same time as those for the Addiction cohort. Mice in the 1-Day Cohort received a single subcutaneous injection of saline or 50mg/kg morphine sulfate and were sacrificed 2 hours later for slice electrophysiology.

**Figure 1:**
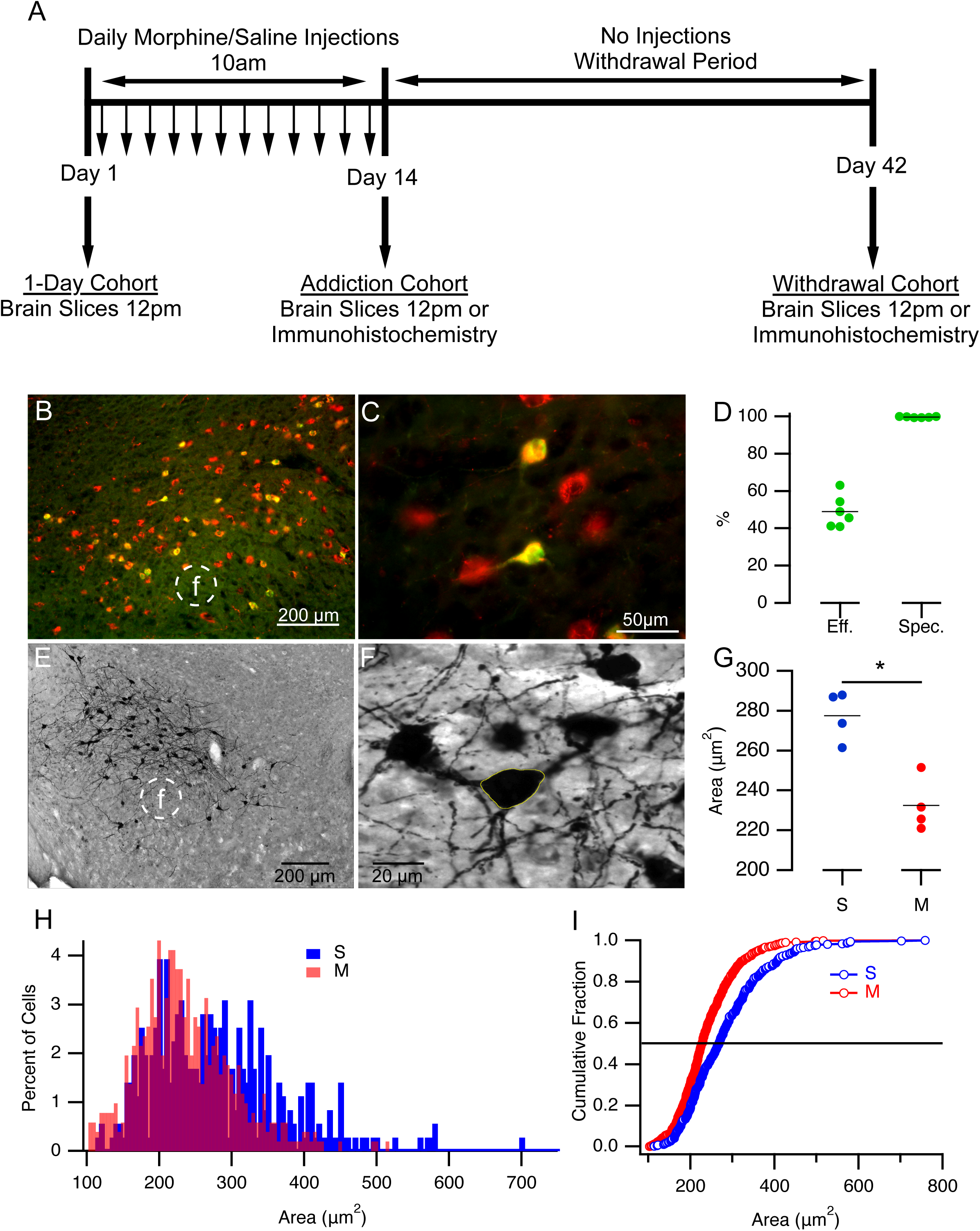
Orexin-EGFP soma size was decreased following chronic morphine. **A.** Schematic of experimental cohorts, injection timing and tissue preparation. **B.** Low power (4x objective) and **C.** higher power (40x objective) epifluorescence images of EGFP-positive neurons within the orexin neuron field. Anti-orexin-A immunofluorescence (red), anti-GFP immunofluorescence (green). **D.** Efficacy (percent orexin immunopositive neurons also expressing GFP immunofluorescence) and specificity (percent GFP immunopositive neurons also expressing orexin immunofluorescence). **E**., **F**. Brightfield low power (E) and higher power (F) of GFP-immunopositive neurons visualized with VIP. Yellow polygon delineates the soma outline (F) used for measurements in G-J. **G**. GFP-immunopositive soma area per mouse from saline-treated (S; blue symbols) and morphine-treated (M; red symbols) mice. **H.** Histogram of all measured GFP-immunopositive soma sizes from each treatment group (S, blue, n = 356 cells; M, red, n = 440 cells). **I**. Cumulative fraction of soma sizes from each treatment group (S, blue symbols; M, red symbols; Kolmogorov-Smirnov test p < 0.0001. Abbreviations: f, fornix; S, saline-treated; M, morphine-treated. Numbers are reported as mean ± sem.

### Brain slice preparations and in vitro electrophysiological recordings

Following deep anesthesia with isoflurane (>5% in O_2_), mice were decapitated, and their brains were quickly removed into ice-cold artificial cerebrospinal fluid (ACSF) which contained (in mM) 126 NaCl, 3 KCl, 1.2 NaH_2_PO_4_, 2 CaCl_2_, 2 MgSO_4_, 24 NaHCO_3_, 10 D-Glucose, osmolarity 295-300, and was oxygenated with carbogen (95% O_2_ and 5% CO_2_). The brain was blocked by first resting the brain on its ventral surface and making coronal cuts caudally through the midbrain-pons junction and rostrally just behind the NAc to leave a block containing the hypothalamus. This block was then affixed with cyanoacrylate to the stage of a Leica VT1000S vibratome, caudal side down with the ventral side of the brain facing the blade. Brain slices (250 µm) were then cut in oxygenated ice cold ACSF and then incubated at room temperature in oxygenated NMDG recovery solution containing (in mM) 115 NMDG, 2.5 KCl, 1.2 NaH_2_PO_4_, 0.5 CaCl_2_, 10 MgSO_4_, 25 NaHCO_3_, 25 D-Glucose, osmolarity 295-300, for 10 minutes (Ting et al., 2014). Slices were then rinsed five times in ACSF and transferred to continuously oxygenated, room temperature ACSF for one hour before starting recordings.

We recorded EGFP-positive orexin neurons in brain slices submerged and perfused (1-2 ml/min) with room temperature (22° ± 2°C) ACSF in a chamber mounted on a fixed stage microscope (Olympus BX-50WI). EGFP positive neurons were selected for recording using epi-fluorescence illumination with a white LED light source (UHP-T-W50-SR; Prizmatix, Holon, Israel) filtered with an EGFP filter cube (#49002; Chroma, VT, USA) and imaged with a CCD camera (Qimaging Retiga Electro; Teledyne Photometrics, AZ, USA). Patching the selected neuron was guided using infrared differential interference contrast (IR-DIC) optics with the same camera operated with Ocular software (Teledyne Photometrics, AZ, USA). We recorded orexin neurons in whole-cell voltage and current clamp (I-clamp fast) configurations using an Axopatch 200B amplifier (Molecular Devices; CA, USA) with the output filter set at 5 KHz. Membrane currents and voltages were sampled at 10 KHz and controlled using a Digidata 1550B and pCLAMP11 software (Molecular devices; CA, USA). Recording pipettes (4-7 Mohm; Sutter Instruments Item # BF150-86-10HP) were made using a horizontal puller (model P-97, Sutter Instruments; CA, USA). Pipettes were filled with an internal solution containing (in mM) 144 K- Gluconate, 0.2 EGTA, 3 MgCl_2_, 10 HEPES, 0.3 NaGTP, 4 Na_2_ATP of pH 7.2 and osmolarity 295-305. Alexa Fluor 568 biocytin or Alexa Fluor 594 biocytin (0.2 mg/ml Thermofisher Scientific) was added to the patch solution to identify the recorded cell and confirm that it was EGFP-positive.

Formation of a Giga seal in voltage clamp mode was monitored using the Membrane Test routine in Clampex 11. Upon seal formation (> 1 Gohms), the fast and slow pipette capacitance compensation was adjusted to minimize stray pipette capacitance. Following breakthrough, the Membrane test routine was used to estimate and monitor series (access) resistance (Ra), total capacitance (C_m_) and input resistance (R_m_) in response to a 50 ms duration −10 mV voltage jump from a holding potential of −60 mV. These values were determined by Clampex 11 from real-time exponential fitting of the average transient current decay (n = 20) and a measure of the average baseline and steady-state current following the decay as described in the Membrane Test Algorithms support page (https://support.moleculardevices.com/s/article/Membrane-Test-Algorithms). Ra was tracked for three minutes to be sure it remained low and stable (17.1 ± 1.0 Mohms, n = 68). If it was unstable or exceeded 40 Mohms the recording was terminated. In principle, these values of Ra should not affect our estimates of R_m_ and C_m_ since this algorithm estimates C_m_ from the total charge delivered during the capacitive transient. Delivery of charge was not limited by Ra or the amplifier since peak currents were less than 2 nA and the Axopatch 200B is capable of sourcing 20 nA. If Ra, C_m_ and R_m_ were stable, we sampled five times over this period, and the average of these values was used as the estimate of these parameters for that cell. Since specific membrane capacitance (∼1 uF/cm^2^) is considered a constant (Gentet, Stuart and Clements, 2000), membrane capacitance is proportional to membrane surface area. Capacitance estimated by this method is most influenced by somatic and proximal dendritic membrane (Taylor, 2012) since these compartments experience most of the voltage change produced by our clamp step.

The holding current was then recorded at −60 mV for 30s and the amplifier was put in the I = 0 mode to measure the resting membrane potential or spontaneous firing of the neuron. The amplifier was then switched into I clamp fast mode to execute a series of current clamp protocols. Series resistance was generally uncompensated unless it was greater than 20 Mohms, since the voltage error from our injected currents were small.

Mouse orexin neurons can be identified as either D-type or H-type based on their membrane potential trajectory following strong hyperpolarizing current pulses delivered while the neurons were spontaneously firing. D-cells show a depolarizing recovery response and rapid resumption of spiking while H-cells show a delayed recovery produced by an A-current mediated hyperpolarization that delayed resumption of spiking (Williams et al., 2008b; Schöne et al., 2011). We therefore identified whether the recorded cell was D- or H-type by delivering a series of increasing amplitude, one second duration current pulses from baseline firing. We then computed a “Spike Ratio” and measured the time to first spike after the end of the pulse, to compare with previously identified D- and H-cells (Schöne et al., 2011). Time to the first spike was measured starting 50 ms after pulse offset to the time of the following spike. “Spike Ratio” was measured as the number of spikes in the one second starting 50ms after the end of the pulse, divided by the number of spikes in the one second before the pulse. (see Fig 2B, C).

**Figure 2:**
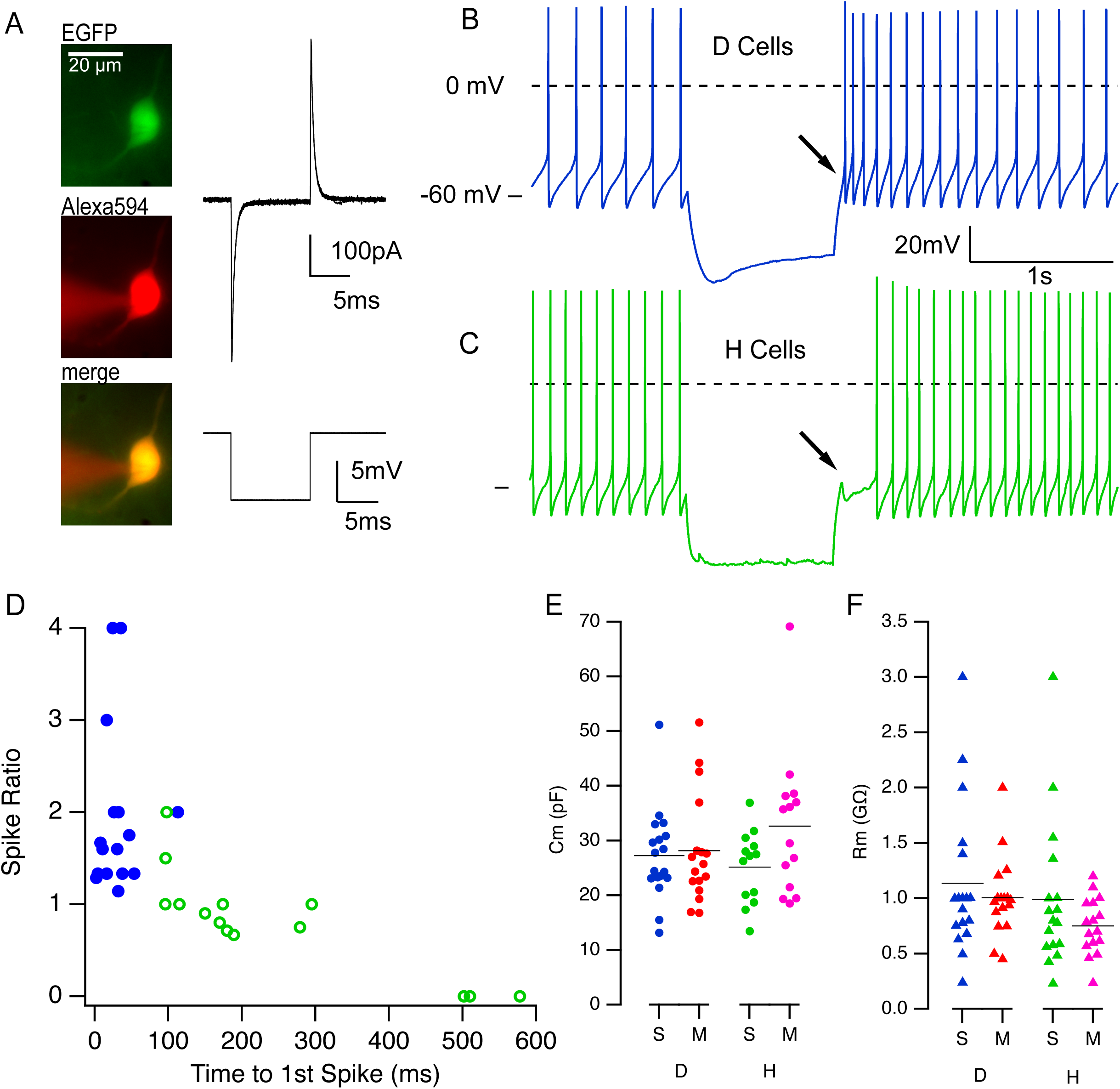
Neither capacitance nor input resistance was altered by chronic morphine in D- or H-type orexin neurons. **A.** Recorded orexin neuron visualized by fluorescence from expressed EGFP (left, top) and Alexa-594- biocytin in the pipette solution (left, middle; images merge left, bottom). Corresponding membrane current (right, top) and voltage (right, bottom) traces from this recorded cell used to estimate C_m_ and R_m_. **B**., **C.** Example voltage traces from a D-cell (B) and H-cell (C). Arrows point to the characteristic depolarizing (D-cells) or hyperpolarizing (H-cells) membrane responses following termination of a hyperpolarizing current pulse. **D.** Spike ratio versus time to first spike following the hyperpolarizing current pulse used to elicit D and H-type responses. **E.** Distribution of C_m_ for D-cells (D) and H-cells (H). C_m_ from D-cells in slices from saline- (blue symbols) or morphine- (red symbols) treated mice, or from H- cells in slices from saline- (green symbols) or morphine- (pink symbols) -treated mice. **F**. Distributions of R_m_ for D- and H- cells from Saline- and Morphine - treated mice (colors as in E).

We next determined the action potential properties by measuring the rheobase and the shape of single spikes. Rheobase was measured as the amount of current required to elicit a single action potential from a holding potential of −60 mV using current pulses of 80 ms duration. Action potential shape was measured by spikes evoked by a current strength just above rheobase. We then measured the F-I relation with firing evoked by two second duration constant current pulses that were incremented in amplitude in 10 pA steps. Reported membrane potentials were not corrected for the liquid junction potential which we previously measured for these solutions as ∼-15 mV.

### Perfusion & brain section collection for Immunohistochemistry (IHC) or Immunofluorescence (IF)

Mice were overdosed with a ketamine (100 mg/kg)/xylazine (10 mg/kg) cocktail and perfused through the heart with 0.01M phosphate buffered saline (PBS) followed by 4 % PFA, two hours following their last injection. Brains were removed and post-fixed in 4 % PFA for 24 hours, after which they were cryoprotected in 30 % sucrose for at least 72 hours. For sectioning, brains were flash frozen in −80° C methylbutane and then frozen into a block of OCT freezing compound (Fisher HealthCare Tissue Plus Cat# 23-730-571) at −20°C. Brain blocks were then sectioned into 40 µm coronal sections using a Leica Cryostat CM1850. 40 µm sections were collected and stored in 0.01M PBS solution prior to immunohistochemistry or immunofluorescence labeling.

### Anti-Orexin-A and Anti-GFP IHC

Brain sections (40 µm) through the lateral hypothalamic area were taken 160 µm apart (every 4^th^ section), and immunostained to visualize either Orexin-A or GFP. Sections were first incubated for 30 minutes with 1% H_2_O_2_ to block endogenous peroxidase activity and then for 30 minutes in 1 % BSA/0.2 % triton-X 100/0.01 M PBS solution to block nonspecific antibody binding. For identification of orexin-A, sections were incubated in rabbit polyclonal anti-orexin-A (1:10,000, Phoenix Pharmaceuticals, H-003-3 lot#01651-8). For identification of GFP, sections were incubated with sheep polyclonal anti-GFP antibody (1:1,000, Novus Biologicals, #NB100-62622, lot#1710). Sections were incubated in primary antibody solution (1 % BSA/0.2 % triton-X 100/ 0.01 M PBS) at room temperature on a shaker overnight. The following morning, primary antibodies were removed, and sections were washed in 0.01M PBS to remove any remaining antibody. Sections were then incubated with avidin biotin complex from a Vectastain Elite ABC Kit (PK-6100 lot#ZF0802) for 30 minutes. Sections were then washed in 0.01 M PBS and then reacted with Vector VIP peroxidase substrate kit (SK-4600) for enough time to obtain optimal purple staining (2-10 minutes). Sections were then mounted onto charged microscope slides (Fisher Scientific, Superfrost plus) and dehydrated in graded ethanol’s (80 %, 90 %, 100 %) for approximately two minutes each to achieve optimal removal of background VIP and dehydrate the tissue. Sections were then cleared with xylene and cover slipped using Permount mounting media.

### Anti-Orexin-A and Anti-GFP IF

Brains were sectioned as above, and sections were incubated for 30 minutes in 1% BSA/0.2% triton-X 100/0.01 M PBS solution to block nonspecific binding. For identification of orexin neurons via orexin-A and GFP IF, the same primary antibodies, at the same concentrations as above were used. Following incubation with primary antibodies overnight in 1 % BSA/0.2 % triton-X 100/ 0.01 M PBS, antibodies were removed, and sections were washed in 0.01 M PBS. Sections were then incubated for 45 minutes in either 1:200 donkey-anti-rabbit-Alexa 594 or donkey-anti-sheep-Alexa 488 in 1 % BSA/0.2 % triton-X 100/ 0.01 M PBS. Sections were washed in 0.01 M PBS a final time and then sections were mounted on charged microscope slides, and cover slipped with anti-fade Vectashield (H-1700) aqueous mounting media.

### Mapping recorded orexin neurons

Low power digital images of the brain slice containing the fornix, mammillothalamic tract, and patch pipette at the location of the last patched neuron were acquired using a 4x objective following each whole cell recording. These images were imported into the FiJi distribution of ImageJ (Schindelin et al., 2012) calibrated and the distance of the tip of the patch pipette to both the fornix and mammillothalamic tract was measured using the line tool and the ROI manager. Corresponding atlas pages were selected from the Paxinos Mouse atlas (Franklin and Paxinos, 2008) and opened in Adobe Illustrator. The measurements taken in ImageJ were then used to plot the location of each recorded neuron onto the atlas page in Adobe Illustrator. A vertical line was drawn through the center of the fornix to separate the nucleus into medial and lateral regions.

### Experimental Design and Statistical Analysis

All investigators were blinded to the mouse treatment conditions for data collection and analyses.

#### Electrophysiological analysis

Analysis of electrophysiological waveforms was performed using Igor Pro software (version 8 and 9; Wavemetrics, OR, USA). Action potential shape and repetitive firing parameters were measured using custom-programmed routines in Igor Pro (Erisir et al., 1999). Action potentials were evoked from a holding potential of −60 mV by injecting 10 pA current steps until reaching threshold to generate one spike. Spike amplitude was measured off-line as the difference between the action potential peak and its threshold. Spike threshold was determined as the potential at which the second derivative of the voltage waveform exceeded three times its standard deviation in the period preceding spike onset. The afterhyperpolarization (AHP) was measured as the difference between the spike threshold and the voltage minimum following the action potential peak. Maximum rates of rise and decay of the action potential were computed from the maximum and minimum of the smoothed first derivative of the voltage waveform. Spike width was measured at half the spike amplitude.

Instantaneous frequency (1/interspike interval) vs time curves were computed from trains of action potentials evoked by 2s duration current pulses from the times after pulse onset at which each spike crossed 0 mV. Initial firing rate was the instantaneous frequency of the first interval and the final rate or steady-state rate was computed as the average of instantaneous frequency for the last five intervals of a train. Instantaneous frequency for the 1^st^, 2^nd^, 4^th^, and the last intervals along with steady-state firing rate were plotted as a function of the injected current strength to construct firing rate – current (F – I) curves. In the repetitive firing protocol, current strength was increased at 10pA increments until spike failure occurred within the 2s duration pulse. Firing rates between 10 – 120 pA were used to compare F- I relations between cells and across treatment conditions and did not include firing at current strengths that produced spike failures. For analysis of firing rate variability, we computed the coefficient of variation (standard deviation of firing rate / mean firing rate) for the period of firing following early accommodation for each firing rate trace at a mid-current strength (80 pA). To estimate the firing rate gain, average F-I curves were fit with a second order polynomial:

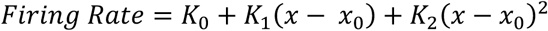

where K_0_ is the firing rate for the minimal current x_0_ (10 pA) and K_1_ and K_2_ are the coefficients derived from the best fit using the built-in curve fitting function of Igor Pro 9 which makes use of a singular value decomposition algorithm. Firing gain was then computed in Igor Pro 9 by taking the 1^st^ derivative of the best-fit polynomial.

#### Histological data collection & Analysis

Every 4^th^ section (40 μm) containing EGFP and orexin-A immunopositive neurons were imaged with a Keyence microscope (BZ-X800) using a 4x (Nikon CFI PlanApo Lambda, NA 0.2) or a 40x (Nikon CFI Plan Apo Lambda, NA 0.95) objective. To prepare images for cell counting, a series images (40x objective) spanning the entire orexin cell field on one side of the brain were acquired and automatically stitched together. The number of Orexin-A or GFP immunopositive neurons was then counted using the ImageJ cell counter plug-in from the stitched images. Neurons were counted from both the left and right sides of the brain, across all sections containing orexin neurons. On average, seven sections per brain (160 µm apart), covered the rostro-caudal extent of the orexin field. Efficacy and specificity of EGFP expression in the orexin-EGFP mice was determined using double immunofluorescence and counting the number of neurons single and double-labelled for orexin-A (Alexa-Fluor 594) and GFP (Alexa-Fluor 488) in untreated mice. Efficacy was calculated as the percent orexin-A positive neurons that were also GFP positive for each mouse. Specificity was measured as the percentage of GFP positive cells that were also orexin-A positive for each mouse.

To estimate efficacy for tissue stained with a single permanent VIP stain for GFP, adjacent sections were stained for orexin-A, and the percent of GFP labeled cells per orexin-labelled cells on the adjacent section was calculated. This allowed us to estimate if morphine treatment grossly changed the efficacy of GFP expression in orexin neurons.

For analysis of soma size, neurons were measured from Z-stacks acquired with the 40x objective using 0.3 µm Z-steps. Z-stack locations were chosen randomly from throughout the GFP+ neuron field. A typical Z stack had 30 images to encompass the entire orexin neuron soma. GFP neurons were only measured if the soma was completely encompassed by the Z-stack, and dendritic/axon projections were visible coming off the soma. Soma measurements were done using the polygon tool in ImageJ to trace the outline of each soma in the z-stack. Measurements of somatic cross-sectional area and other shape parameters were collected in the region of interest manager in ImageJ.

#### Statistical Analysis

Statistical analyses were conducted using DataDesk RP (v8.2.1; Data Descriptions, Ithaca, NY), Excel, or Igor Pro 8 and 9. Data are presented as the mean ± standard error of the mean (SEM). For all statistical tests, p < 0.05, p < 0.01, and p < 0.001 were considered significant, and p > 0.05 was considered not significant (ns). For histological data, average soma size per mouse was compared between treatment groups by T-test, and cumulative distributions of individual cell size were compared with a Kolmogorov-Smirnov test implemented in Igor Pro. For electrophysiological data: analysis of C_m_, R_m_, Rheobase, spontaneous firing and action potential shape parameters were compared between treatments groups with a one-way ANOVA. Analysis of different firing categories (Continuous, Cluster, Reluctant) was done via a Chi-Square test. F – I curves between treatment groups were analyzed using two-way repeated measures ANOVAs (rmANOVA) set up using a general linear model with firing rate as the dependent variable and treatment and current strength (repeat factor) and the interaction between treatment and current strength as the factors. Differences between firing rate at each current strength was determined by Bonferroni corrected post-hoc tests.

## Results

### Chronic morphine exposure reduced average soma size of orexin-EGFP neurons

To study the effects of chronic morphine on excitability, we first confirmed the efficacy and specificity of EGFP expression in orexin neurons and then determined whether EGFP-expressing orexin neurons reflect the decrease in soma size previously reported for the orexin immunoreactive population (Thannickal et al., 2018).

Using double immunofluorescence we found that that efficacy was variable but specificity was very high (Fig 1 B – D). Efficacy ranged between 40 – 70 % across animals with an average of 49.1 ± 3.5 % (n = 6) of orexin-A immunopositive neurons expressing EGFP (Fig 1D, left), while the specificity of orexin-A expression within the GFP positive population was 99.7 ± 0.1 % (n = 6; Fig 1D, right).

We next determined if chronic morphine treatment altered soma size of EGFP-expressing orexin neurons. Male orexin-EGFP mice in our Addiction cohort were injected with 50 mg/kg morphine or saline once daily for two weeks prior to VIP IHC for GFP (Fig 1 E, F). We found that the average soma area was reduced by ∼16 %, decreasing from 277.5 ± 6.2 µm^2^ in saline treated mice (n = 4) to 232.5 ± 6.7 µm^2^ in morphine treated mice (n = 4; Fig 1G). Pooling the soma measurements from all mice in each treatment (Saline treated: n = 356; Morphine-treated: n = 440), indicated a leftward shift in the size distribution for morphine treated mice and suggested that the average size was decreased due to a modest increase in small somata and a large decrease in large somata (Fig 1H). This leftward shift was apparent by comparing the cumulative size distributions which were significantly different based on a Kolmogorov-Smirnov test (p << 0.0001; Fig 1I). Thus, these EGFP+ neurons undergo a similar reduction in soma size to that previously found for the overall population of orexin immunopositive neurons following chronic morphine exposure.

### Chronic morphine exposure did not modify the basic passive electrical properties of D- or H-type orexin neurons

We next assessed the passive electrical properties of orexin-EGFP neurons in brain slices from mice in the Addiction cohort by whole-cell recording (Fig 2A, left). Total membrane capacitance (C_m_) and input resistance (R_m_) were measured from a holding potential of −60 mV using the Clampex membrane test measurements of capacitive currents resulting from −10 mV voltage pulses (Fig 2A, right). Since Orexin-EGFP neurons had smaller somata following 14 days of morphine exposure, we expected that estimates of C_m_ might be lower and R_m_ might be higher following morphine treatment, however this was not the case. There were no significant differences between EGFP-positive cells from saline- and morphine-treated mice in either C_m_ (Saline: 25.64 ± 1.29 pF, n = 35; Morphine 29.87 ± 1.97 pF, n = 33; t = −1.79, df = 55, P = 0.08) or R_m_ (Saline: 1.06 ± 0.11 Gohm, n = 35; 0.88 ± 0.06 Gohm, n = 33; t = 1.44, df = 50, P = 0.16) although there was a trend toward a higher average Cm among neurons from the morphine treated mice.

Orexin neurons are in an intrinsic state of depolarization and typically fire spontaneously in brain slices (Li et al., 2002; Eggermann et al., 2003; Cvetkovic-Lopes et al., 2010). We found that 64/68 neurons fired spontaneously but there was no statistical difference in the average rate between EGFP positive cells from saline (8.55 ± 0.80 Hz, n = 32) and morphine (8.59 ± 0.98 Hz, n = 32; t = 0.172, df = 60, P = 0.98) -treated mice. Moreover, we found no statistical difference in variability of spontaneous firing as measured by the coefficient of variation (CV; Saline: 0.15 ± 0.02, n = 32; Morphine: 0.17 ± 0.02, n = 32; t = −0.697, df = 60, P = 0.49).

We next characterized each neuron as D- or H- subtypes (Williams et al., 2008b; Schöne et al., 2011) by delivering hyperpolarizing current clamp pulses while they were spontaneously firing (Fig 2B, C). Out of 68 cells tested in the Addiction Cohort, 34 were classified as D-type, with a rapid return to firing after offset of the current pulse (arrow) and 28 were classified as H-type, with a delayed return to firing after offset of the current pulse (arrow). Four neurons were not spontaneously firing and were not classified. We also found these orexin neurons showed a similar relation between spike ratio vs. time to first spike, as previously described (Schöne et al., 2011). D-type neurons had higher spike ratios and shorter times to first spike compared to H-type orexin neurons (Fig 2D).

Analyzing C_m_, R_m_ or spontaneous firing rate separately for each cell type also did not reveal an effect of treatment on these parameters, although the trend toward higher C_m_ and lower R_m_ following morphine treatment was manifest only for H-neurons (Fig 2 E, F; Table 1). Thus, despite the soma size reduction after two weeks of daily morphine exposure, electrical measurement did not reveal a corresponding decrease in C_m_ or increase in R_m_. Several factors may contribute to this apparent discrepancy. First, cell sampling differences between fixed sections and brain slices may have masked differences in the size distributions. Second, if soma size alone decreased, the effect on total capacitance would become much less detectable, since soma area is only ∼15 % of total membrane area (Schöne et al., 2011). Finally, small membrane infoldings may contribute to soma size decrease without reducing total membrane area and hence capacitance (Matovic et al., 2020).

**Table 1.** Cell type properties from mice treated for two weeks with saline or morphine. Passive and active properties of D-cells (left columns) and H-cells (right columns) from mice treated for two weeks with a single daily injection of saline or morphine (50 mg/kg). Measured parameters are displayed as mean ± SEM. Columns show D-cell data from mice treated with saline (D-type (Saline)) or morphine (D-type (Morphine)). Each measurement is followed by the number of cells (n) measured. The value in the column labelled P is the probability of rejecting the null hypothesis when it is true that the Saline and Morphine values were drawn from the same distribution computed from a two-tailed, two-sample T-test with unequal variance.

|  | D-type (Saline) |  | D-type (Morphine) |  |  | H-type (Saline) |  | H-type (Morphine) |  |  |
| --- | --- | --- | --- | --- | --- | --- | --- | --- | --- | --- |
|  | Mean ± SEM | n | Mean ± SEM | n | P | Mean ± SEM | n | Mean ± SEM | n | P |
| Passive Properties |  |  |  |  |  |  |  |  |  |  |
| Capacitance (pF) | 27.26 ± 1.96 | 17 | 28.16 ± 2.42 | 17 | 0.775 | 25.17 ± 1.85 | 14 | 32.67 ± 3.55 | 13 | 0.076 |
| Input Resistance (Gohm) | 1.13 ± 0.16 | 17 | 1.01 ± 0.09 | 17 | 0.485 | 0.99 ± 0.17 | 14 | 0.75 ± 0.06 | 13 | 0.186 |
| Active Properties |  |  |  |  |  |  |  |  |  |  |
| Rheobase (pA) | 15.88 ± 1.50 | 17 | 18.24 ± 1.76 | 17 | 0.317 | 22.86 ± 3.22 | 14 | 33.33 ± 3.29 | 13 | <0.05 |
| Spontaneous Firing Rate (Hz) | 8.34 ± 1.02 | 16 | 9.41 ± 1.39 | 17 | 0.544 | 8.77 ± 1.28 | 15 | 7.73 ± 1.51 | 13 | 0.601 |
| Threshold (mV) | -36.97 ± 0.59 | 16 | -37.64 ± 0.66 | 17 | 0.456 | -38.19 ± 0.48 | 15 | -36.70 ± 0.73 | 12 | 0.104 |
| Rising Slope Max (mV/ms) | 203.11 ± 8.06 | 16 | 198.08 ± 10.3 | 17 | 0.703 | 245.59 ± 14.53 | 15 | 203.22 ± 10.10 | 12 | <0.05 |
| AP Max (mV) | 51.35 ± 1.01 | 16 | 49.67 ± 1.32 | 17 | 0.319 | 52.87 ± 1.78 | 15 | 46.73 ± 1.67 | 12 | <0.05 |
| AP Amplitude (mV) | 88.32 ± 1.22 | 16 | 87.31 ± 1.34 | 17 | 0.581 | 91.06 ± 2.08 | 15 | 83.43 ± 1.65 | 12 | <0.01 |
| Falling Slope Max (mV/ms) | -88.91 ± 3.64 | 16 | -87.25 ± 4.49 | 17 | 0.777 | -102.08 ± 5.88 | 15 | -82.15 ± 4.57 | 12 | <0.05 |
| Width at Half Max (ms) | 1.11 ± 0.02 | 16 | 1.14 ± 0.03 | 17 | 0.454 | 1.03 ± 0.03 | 15 | 1.13 ± 0.04 | 12 | <0.01 |
| AHP Min (mV) | -67.01 ± 0.92 | 16 | -67.14 ± 0.75 | 17 | 0.916 | -68.31 ± 0.80 | 15 | -64.89 ± 0.83 | 12 | <0.05 |

### Chronic morphine exposure altered the action potential shape of H- but not D-type orexin neurons

We next assessed the consequences of 14 days of morphine exposure on the active membrane properties underlying spike generation in orexin neurons by measuring rheobase and action potential shape parameters. Orexin neurons from morphine-treated mice had on average, a higher rheobase (25.81 ± 2.40 pA, n = 31) than those from saline-treated mice (19.03 ± 1.76 pA, n = 31; t = 24.178, df = 60, P = 0.026). Analyzing D- and H-cells separately, revealed that this difference was accounted for by the H-cells, which showed a 30% higher average rheobase in slices from morphine-treated mice compared to those from saline-treated mice, while no difference was found among D-cells (Table 1). Thus, daily morphine exposure selectively increases the amount of current required to elicit an action potential in H-cells.

We next analyzed the shape of single spikes produced by just-suprathreshold current injections from a membrane potential of −60 mV (Fig 3, Table 1). This revealed that the action potentials in H-cells from morphine-treated mice were, on average, smaller and slower than action potentials from saline-treated mice (Fig 3A, right). Moreover, there were no corresponding changes to the action potential shape in D-cells recorded from the same brain slices (Fig 3A, left). In addition to having action potentials of lower amplitude (Fig 3B) and longer half-widths (Table 1), morphine treated H-cells had action potentials with significantly slower maximal rising (Fig 3D, Table 1) and falling rates (Fig 3E, Table 1) with a significantly less negative average peak afterhyperpolarization (Fig 3C, Table 1), while there were no corresponding differences between these parameters for D-cells (Table 1).

**Figure 3:**
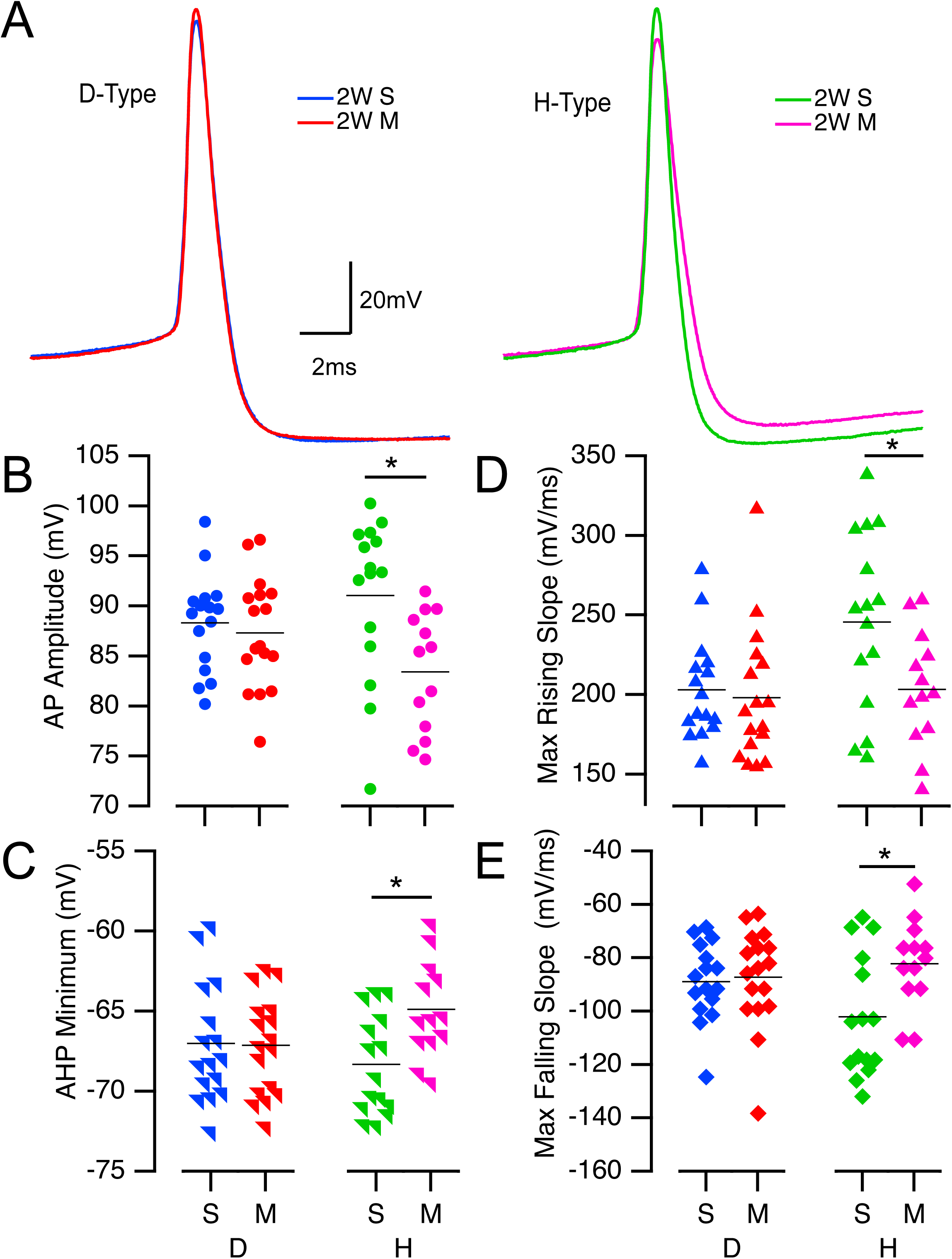
H-type, but not D-type orexin neurons have smaller, wider action potentials following two weeks of chronic morphine. **A.** Superimposed action potentials from representative D-cells following saline- (blue) or morphine- (red) treatment (left) and from H-cells following saline- (green) or morphine- (pink) treatment (right). **B – E**. Distributions of spike shape parameters for D-cells (left) and H-cells (right) following either saline (S) or morphine (M) treatment (colors as in A). The average spike amplitude (B; AP Amplitude), minimum value of the afterhyperpolarization (C; AHP Minimum), maximum rate of rise (D; Max rising slope) and maximal falling rate (E; Max Falling Slope) were all altered for H-cells, but not D-cells from morphine-treated mice compared to saline-treated mice. B – E. Horizontal lines indicate mean values.

Despite these changes in spike shape, there were no indications that H-cells from morphine-treated mice were more depolarized than their counterparts from saline-treated mice. Neither their spontaneous firing rates were different (Table 1), nor was a greater negative current required to hold H-cells from the morphine-treated mice at −60 mV (holding current from saline-treated = −24.9 ± 5.2 pA, N = 15; from morphine-treated = −30.1 ± 10.4 pA, n = 13; t = 0.61, df = 24, P = 0.65). These data indicate that 14 days of daily morphine exposure results in a selective impairment in the spike generating mechanisms of H-cells while sparing D-cells.

### Chronic morphine exposure impaired repetitive firing in H-type but not D-type orexin neurons

We next examined the ability of orexin neurons to fire repetitively in response to constant current pulses. During this protocol, steady current was injected to hold the baseline membrane potential at −60 mV and two second duration positive current pulses from 10 pA up to 120 pA in 10 pA steps were injected with enough time between each pulse to allow the membrane potential to recover to baseline. Across the injected current steps, we noticed that cells displayed different firing patterns that were reflected in their spike-rate variability. Three patterns were discerned: The largest proportion of cells fired continuously with a regular pattern throughout the current pulse (termed “Continuous”), a smaller portion fired irregularly with pauses in spiking punctuating the train, which we termed “Cluster firing”, and some cells we termed “Reluctant”, only fired one or a few spikes no matter how much current was delivered (Fig 4A). To capture the variability differences between Continuous and Cluster cells, we measured the coefficient of variation (CV) of the instantaneous firing after early adaptation at a mid-current strength of 80 pA for cells in each treatment group (Fig 4B). Based on the distribution of CVs across groups, we chose a CV of 0.22 to separate cells into Continuous (< 0.22) and Cluster (> 0.22) categories. Using this criterion, we observed D-cells to have a high percentage of Continuous firing cells with only a few neurons showing Cluster or Reluctant firing patterns. This distribution didn’t change following morphine exposure (x^2^ = 3.15; df = 2; n = 30; P = 0.248; Fig 4C left). However, assessing the firing patterns of H-type cells revealed that those from morphine treated mice had significantly more Cluster and Reluctant firing cells than H cells from saline treated mice (x^2^ = 9.45; df = 2; n = 24; P = 0.0033; Fig 4C right). CV was inversely related to average firing rate, with H-cells from mice treated with morphine comprising most of the cells with high CVs and low firing rates for the same current strength (Fig 4D). These data indicate that 14 days of daily morphine increases the variability in firing of H-cells but not D-cells and may impair the ability of H-cells to repetitively fire.

**Figure 4:**
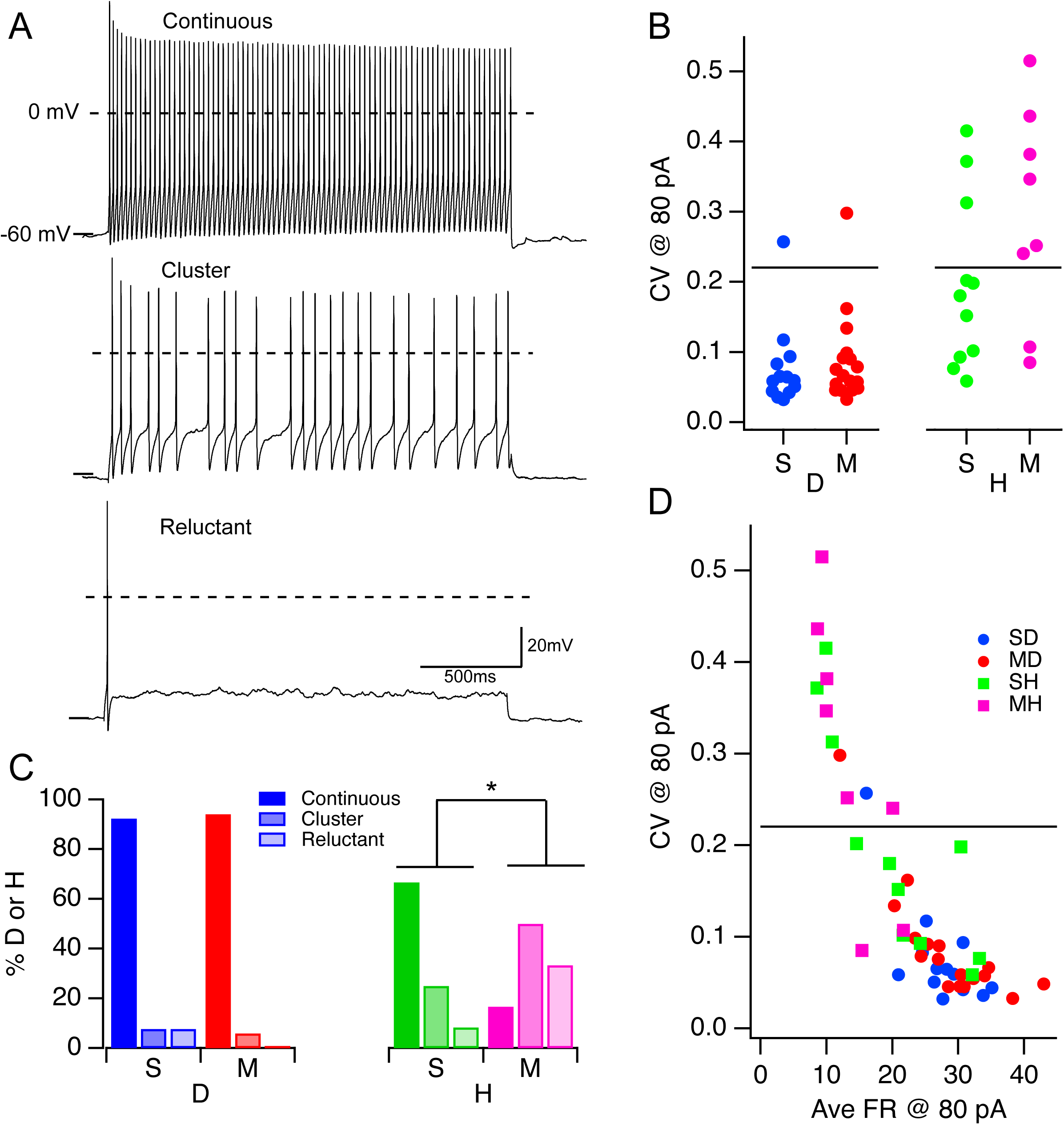
Chronic morphine increased spike-rate variability in H-type, but not D-type orexin neurons. **A.** Repetitive firing examples from three orexin neurons in response to mid-range depolarizing currents illustrating “Continuous” (top), “Cluster” (middle), and “Reluctant” (bottom) firing patterns. **B**. Distribution of firing rate coefficient of variations (CVs) resulting from an 80 pA current step for D-cells (left) from saline (S; blue symbols) and morphine (M; red symbols) -treated mice and H-cells (right) from saline (S; green symbols) and morphine (M; pink symbols) -treated mice. A CV of 0.22 at 80 pA of current injection (horizontal lines) was selected to distinguish continuous (< 0.22) and cluster (> 0.22) firing types. **C**. Percent of D- (left) and H- (right) cells identified with each firing pattern. Colors indicate cell-type and treatment group as in B. Firing pattern indicated by bar saturation (Continuous: 100%; Cluster: 50% and Reluctant: 25%) **D**. Coefficient of variation (CV @ 80 pA) versus average firing rate at 80 pA of current injection (Ave FR @ 80 pA), horizontal line indicates a CV of 0.22. Colors indicate cell-type and treatment group as in B.

To examine repetitive firing ability, we compared the initial and final firing rates achieved in response to two second current pulses from 10 – 120 pA in cells from mice treated with either saline or morphine. Sample D-cell recordings are shown in Fig 5A (saline-treated left, morphine-treated right) and sample H-cell recordings are shown in Fig 6A (saline-treated left, morphine-treated right). We computed instantaneous firing vs. time curves for each current strength (D-cells: Fig 5B; H-cells: Fig 6B) and then constructed average F - I curves. Since Continuous and Cluster firing neurons showed spike frequency adaptation (SFA), we constructed F - I curves for both the initial firing rate, measured between the first two spikes, and the final firing rate, measured as the average rate of the last seven spikes, or the average over two seconds when there were fewer than seven spikes. F - I curves were then averaged across cells and the average curves (± SEM) for D-cells and H-cells from each treatment condition are illustrated in Fig 5C and 6C, respectively.

**Figure 5:**
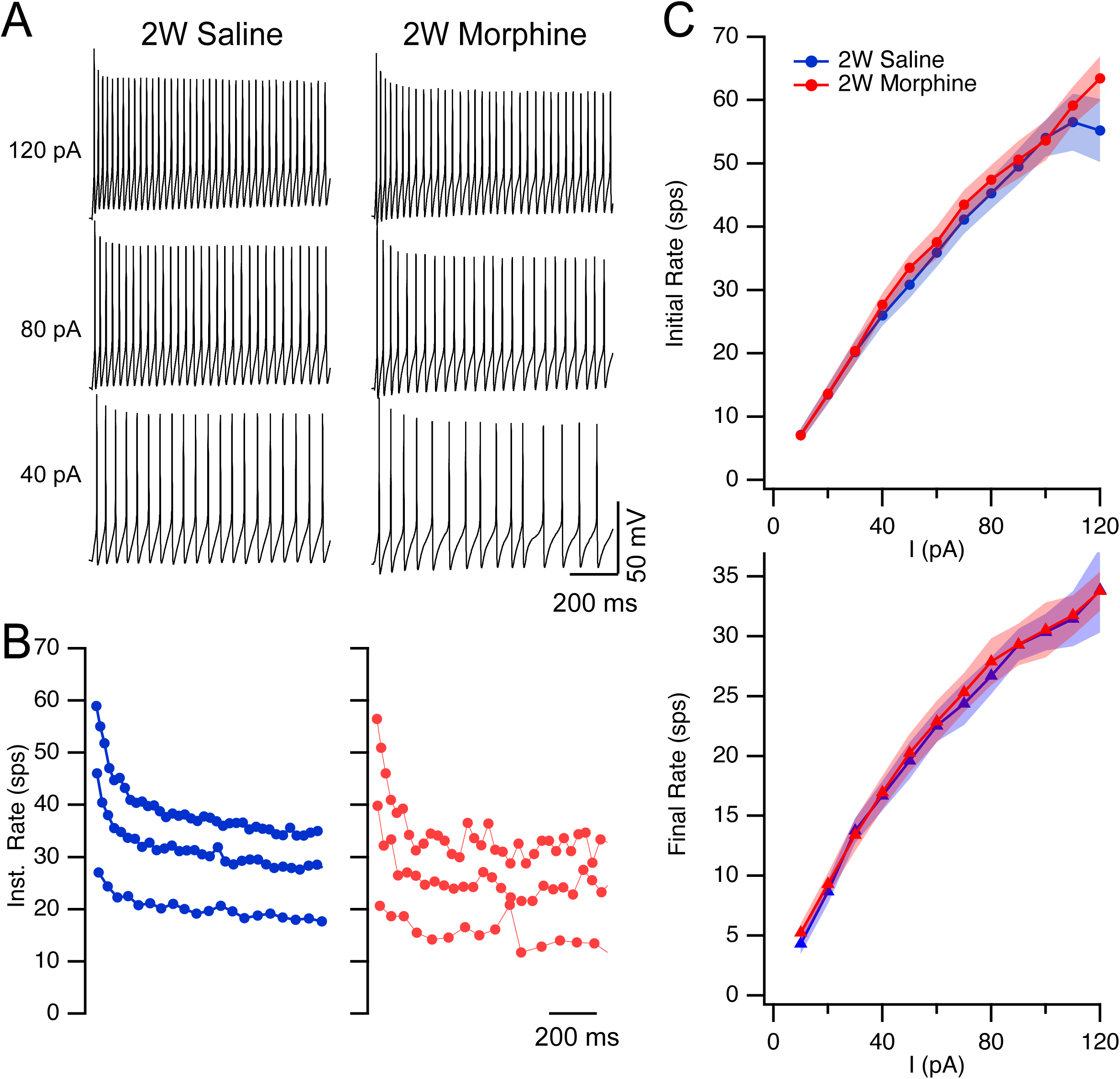
Chronic morphine exposure did not alter the firing capability of D-type orexin neurons. **A.** Example repetitive firing of D-cells with continuous firing patterns evoked by 40, 80, and 120 pA current pulses from a saline- (left) and a morphine-treated (right) mouse. **B.** Corresponding instantaneous firing rate curves for the traces in A (saline: blue symbols; morphine: red symbols). **C**. Initial Rate (Mean ± SEM) vs. current (I) curves (Top) and Final Rate (Mean ± SEM) vs. current (I) curves (Bottom) for D-cells. Colors as indicated in B.

**Figure 6:**
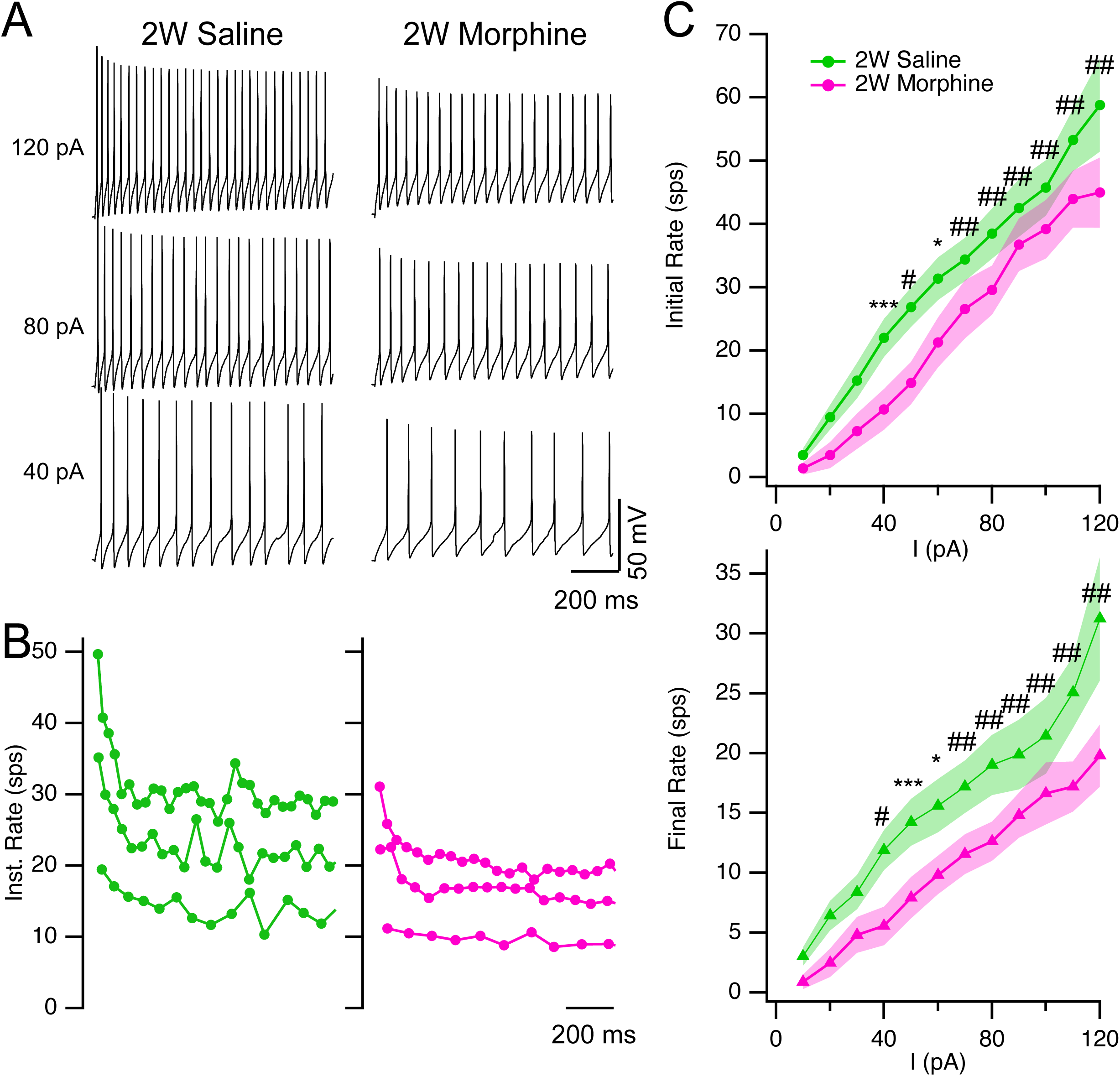
Chronic morphine exposure impaired the firing capability of H-type orexin neurons. **A**. Example repetitive firing of H-cells with continuous firing patterns evoked by 40, 80, and 120 pA current pulses from a saline- (left) and a morphine-treated (right) mouse. **B**. Corresponding instantaneous firing rate curves for the traces in A (saline: green symbols; morphine: pink symbols). **C**. Initial Rate (Mean ± SEM) vs. current (I) curves (Top) and Final Rate (Mean ± SEM) vs. current (I) curves (Bottom) for H-cells. Colors as indicated in B. Differences between treatments were found for firing at most current strengths as indicated by Bonferroni post-hoc testing. * p < 0.05; *** p < 0.005; # p < 0.001; ## p < 0.0001.

To compare the F - I curves we used a two-way rmANOVAs. Examining the initial firing rate for D-type cells we found that the rmANOVA was not significant for treatment (df = 1, F-ratio = 0.00013, P = 0.99) or for the interaction between treatment and current strength (df = 11, F-ratio = 0.16, P = 1.0) and there were no differences between treatments at any current strength as determined by Bonferroni post-hoc testing (all p values > 0.97; Fig 5C top). Similarly, for D-cell final firing rate, the rmANOVA was not significant for treatment (df = 1, F-ratio = 0.47, P = 0.50) or for the interaction between treatment and current strength (df = 11, F-ratio = 0.59, P = 0.83) nor were there any significant differences between treatments at any current strength as determined by Bonferroni post-hoc testing (all p values = 1.0; Fig 5C, bottom). Thus, two weeks of daily morphine treatment did not alter the repetitive firing ability of D-cells compared to two weeks of daily saline treatment.

In contrast, the average F - I curves for initial and final firing rates for H-cells from mice treated with morphine were largely not overlapping with the F - I curves obtained from mice treated with saline. H-cells from morphine treated mice had on average a 35.0 ± 5.3 % and 40.9 ± 4.1 % decrease in average firing rate across 10-120 pA of current injection for the initial and final firing rates respectively (Fig 6 C). While the rmANOVA for initial firing rate was not significant for treatment (df = 1, F-ratio = 2.36, P = 0.14) or interaction (df = 11, F-ratio = 2.25, P = 0.05), there were significant differences between treatments at most current strengths as indicated by Bonferroni post-hoc testing (Fig 6 C, top). A similar pattern was apparent for the final firing rates. While the rmANOVA for final firing rate was not significant for treatment (df = 1, F-ratio = 1.91, P = 0.19) or interaction (df = 11, F-ratio = 1.96, P = 0.09), there were significant differences between treatments at most current strengths as indicated by Bonferroni post-hoc testing (Fig 6C, bottom). Thus, chronic morphine altered H-cell firing by increasing the numbers of H-cells with Cluster and Reluctant firing patterns and diminishing the ability of continuous and cluster firing H-cells to repetitively fire, while having no detectable effects on the firing ability of D-cells recorded from the same brain slices.

It was also apparent from these recordings that both D- and H- cells showed changes in spike shape during repetitive firing. To determine if our morphine treatment impacted these changes, we measured how spike amplitude, maximal rising rate and halfwidth varied with spike number. Since firing rate can influence these shape changes, successive spike shapes were evaluated from traces where the final firing rate reached ∼10 spikes per second. This revealed that successive action potentials from both D- and H- cells decreased in amplitude, decreased in maximal rising rate and increased in half-width. We found that between the first and tenth spike, action potentials from D-cells recorded in slices from saline-treated mice (n = 13) decreased in amplitude by 25.6 ± 2.3 %, while maximal rising rate decreased by 53.7 ± 3.6 % and half-width increased by 44.5 ± 4.2 %. None of these changes were statistically different in D-cells recorded in slices from mice treated with morphine (n = 17). These action potentials decreased in amplitude by 24.5 ± 1.1 % (df = 28, t = 0.462, P = 0.65), maximal rising rate decreased by 54.8 ± 1.4 % (df = 28, t = 0.308, P = 0.76) and half-width increased by 46.0 ± 3.7 % (df = 28, t = 0.2668, P = 0.79). Over the first 10 spikes, action potentials from H-cells recorded in slices from saline-treated mice (n = 11) decreased in amplitude by 20.8 ± 2.0 %, while maximal rising rate decreased by 47.1 ± 3.6 % and half-width increased by 35.3 ± 4.8 %. In contrast to D-cells, H-cells recorded in slices from mice treated with morphine (n = 8) had action potentials that showed statistically greater changes in these parameters. Action potential amplitude decreased by 30.0 ± 3.9 % (df = 17, t = 2.26, P = 0.04), the maximal rising rate decreased by 59.7 ± 5.1 % (df = 17, t = 2.09, P = 0.05) and the half-width increased by 56.0 ± 8.5 % (df = 17, t = 2.26, P = 0.03). Thus, fourteen daily injections of morphine exaggerated the processes underlying spike amplitude decrease and spike slowing during repetitive firing in H-cells but not in D-cells.

### Four-weeks of withdrawal was sufficient to reverse the soma size decrease of orexin neurons and impaired excitability of H-cells

The decrease in orexin neuron soma size produced by two weeks of morphine exposure was shown previously to recover by four weeks of withdrawal (Thannickal et al., 2018). We therefore determined whether four weeks of withdrawal was sufficient for soma size in the orexin-EGFP subpopulation to recover, and if the impaired excitability of H-cells from morphine-treated mice persisted or recovered.

We measured soma size of orexin-EGFP neurons from mice in the Withdrawal cohort and found that the per-animal soma size averages were not different between groups four weeks after the end of the saline or morphine treatments (Fig 7A; Saline: 191.3 ± 12.89 µm^2^, n = 5; Morphine: 195.56 ± 5.16 µm^2^, n = 5; t = 0.582, df = 8 p = 0.771,). The similarity of cell sizes for the two treatment groups after four weeks of withdrawal was also clearly reflected in the overlapping distributions of individual cell sizes (Fig 7B) which were not statistically different based on the Kolmogorov-Smirnov test (D = 0.007, P = 1.0).

**Figure 7:**
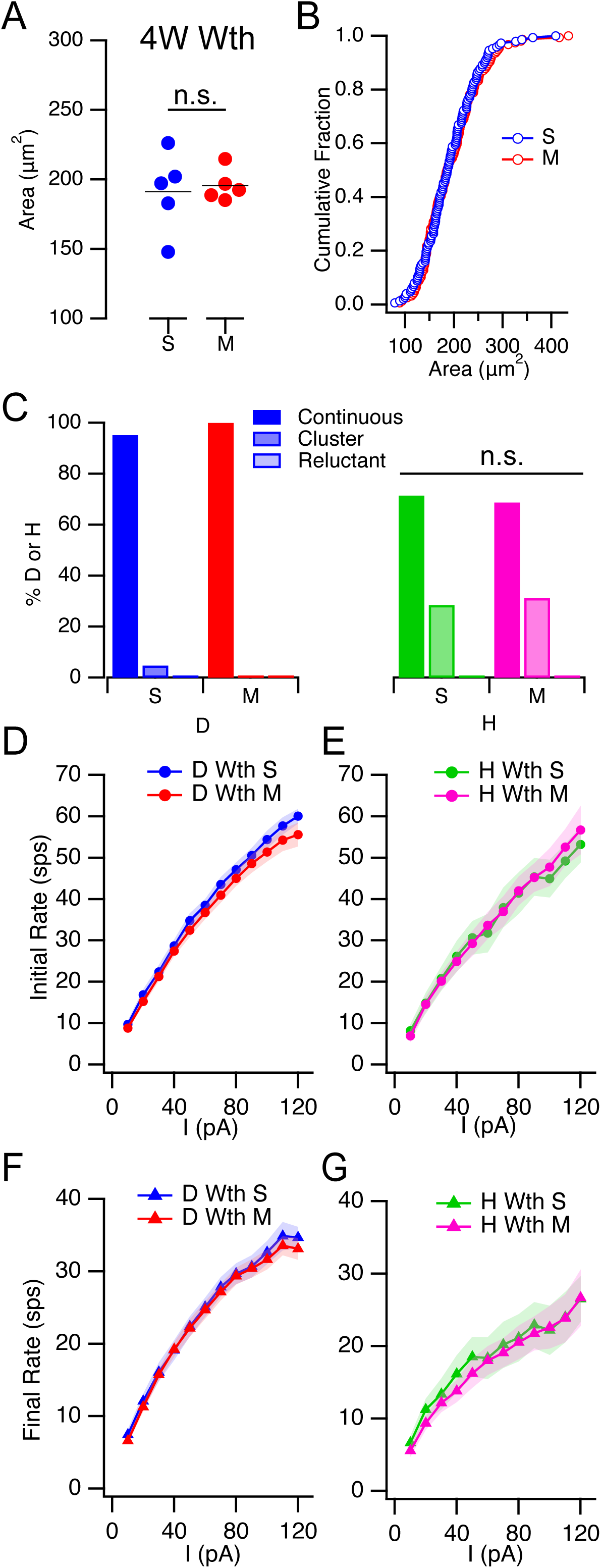
After four weeks of recovery, differences in soma size and intrinsic excitability were no longer apparent. **A**. Average soma size per mouse for saline-treated (S, blue symbols, n = 5) and morphine-treated mice (M, red symbols, n = 5) were not significantly (n.s) different four weeks after the end of treatment. **B.** Cumulative distribution of individual soma sizes from saline-treated (S, open blue symbols, n = 148) and morphine-treated mice (M, open red symbols, n = 156) measured four weeks after the end of treatment. **C.** Distribution of recorded D- (Left) and H-cells (Right) with “continuous”, “cluster” and “reluctant” firing pattern from slices made following four weeks of withdrawal after saline-treatment (S, Blue for D-cells; Green for H-cells) and morphine-treatment (M, red for D-cells; pink for H-cells). The distribution of H-cells from morphine-treated mice was not significantly (n.s.) different from that from saline-treated mice. **D-G.** Firing frequency (mean ± SEM) – current (I) curves for initial rates (D, E) and final rates (F, G) for D- cells (D, F) and H-cells (E, G). Colors as in C. Abbreviations: n. s., not significant; S, saline-treated; M, morphine-treated; Wth, Withdrawal cohort; sps, spikes/second.

Comparing the electrical properties of D- and H-cells four weeks after the end of saline or morphine treatment revealed that H-cell spike parameters that differed between morphine and saline-treated mice in the Addiction cohort were no longer different among H-cells from the Withdrawal cohort (Table 2). Moreover, passive properties and spike-shape parameters that were not different in neurons from the Addiction cohort, were also not different in neurons from the Withdrawal cohort suggesting that no additional changes emerged during this period. Thus, following four weeks of withdrawal, the rheobase observed for H-cells from morphine-treated mice was no longer larger than that for H-cells from saline-treated mice. Similarly, spike parameters like amplitude, maximal rate of rise, maximal falling rate, width at half max, and AHP minimum in morphine treated H-cells were no longer different from those measured in saline-treated H-cells following four weeks of withdrawal. This indicates that the impaired spike-generation mechanisms which resulted in smaller and slower spikes in H-cells following morphine treatment, either recovered or were compensated for by alternative mechanisms following four weeks of withdrawal.

**Table 2.** Cell type properties from mice after four weeks of passive withdrawal. Passive and active properties of D-cells (left columns) and H-cells (right columns) from mice treated for two weeks with a single daily injection of saline or morphine (50 mg/kg) and then allowed to recover for 4 weeks. Measured parameters are displayed as mean ± SEM. Columns show D-cell data from mice treated with saline (D-type (Saline)) or morphine (D-type (Morphine)). Each measurement is followed by the number of cells (n) measured. The value in the column labelled P is the probability of rejecting the null hypothesis when it is true that the Saline and Morphine values were drawn from the same distribution computed from a two-tailed, two-sample T-test with unequal variance.

|  | D-type (Saline) |  | D-type (Morphine) |  |  | H-type (Saline) |  | H-type (Morphine) |  |  |
| --- | --- | --- | --- | --- | --- | --- | --- | --- | --- | --- |
|  | Mean ± SEM | n | Mean ± SEM | n | P | Mean ± SEM | n | Mean ± SEM | n | P |
| Passive Properties |  |  |  |  |  |  |  |  |  |  |
| Capacitance (pF) | 25.31 ± 2.26 | 21 | 32.84 ± 3.11 | 15 | 0.061 | 28.06 ± 2.17 | 14 | 26.74 ± 2.85 | 14 | 0.642 |
| Input Resistance (Gohm) | 1.41 ± 0.18 | 21 | 1.82 ± 0.31 | 15 | 0.252 | 1.46 ± 0.33 | 14 | 1.18 ± 0.19 | 14 | 0.466 |
| Active Properties |  |  |  |  |  |  |  |  |  |  |
| Rheobase (pA) | 11.71 ± 1.27 | 21 | 11.9 ± 1.11 | 20 | 0.913 | 14.78 ± 2.56 | 14 | 14.21 ± 2.47 | 14 | 0.964 |
| Spontaneous Firing Rate (Hz) | 7.75 ± 1.14 | 19 | 7.72 ± 1.09 | 19 | 0.986 | 5.48 ± 1.28 | 13 | 8.46 ± 1.33 | 14 | 0.118 |
| Threshold (mV) | -38.98 ± 0.79 | 22 | -37.36 ± 0.68 | 22 | 0.129 | -37.79 ± 0.80 | 14 | -39.67 ± 0.73 | 14 | 0.141 |
| Rising Slope Max (mV/ms) | 180.45 ± 7.70 | 22 | 173.31 ± 6.46 | 22 | 0.486 | 203.53 ± 11.14 | 14 | 202.46 ± 9.08 | 14 | 0.942 |
| AP Max (mV) | 49.16 ± 1.35 | 22 | 48.78 ± 0.66 | 22 | 0.802 | 51.17 ± 1.26 | 14 | 50.22 ± 1.49 | 14 | 0.632 |
| AP Amplitude (mV) | 88.15 ± 1.46 | 22 | 86.14 ± 0.77 | 22 | 0.235 | 88.96 ± 1.46 | 14 | 86.43 ± 1.62 | 14 | 0.751 |
| Falling Slope Max (mV/ms) | -80.02 ± 3.23 | 22 | -78.22 ± 2.49 | 22 | 0.665 | -89.92 ± 5.38 | 14 | -89.24 ± 4.69 | 14 | 0.925 |
| Width at Half Max (ms) | 1.26 ± 0.04 | 22 | 1.26 ± 0.03 | 22 | 0.958 | 1.17 ± 0.06 | 14 | 1.17 ± 0.03 | 14 | 0.957 |
| AHP Min (mV) | -68.11 ± 0.99 | 22 | -65.80 ± 0.60 | 22 | 0.056 | -67.90 ± 1.42 | 14 | -68.16 ± 1.21 | 14 | 0.893 |

A similar picture emerged from analysis of the repetitive firing ability of D- and H-cells from mice in the withdrawal cohort. After four weeks of withdrawal, H-cells from morphine-treated mice no longer showed an increase in the fraction of cells with cluster or reluctant firing patterns compared to H-cells from saline-treated mice (Fig 7 C; X^2^ = 1.32, df = 2, P = 0.294). Moreover, neither the initial (Fig D, E) nor final F - I relations (Fig 7 F, G) of D- and H-cells from morphine-treated mice were different compared to those from saline-treated mice after four weeks of withdrawal, (Fig 7 D-cells: D, F; H-cells: E, G). This was determined using a 2-way rmANOVA analyses for initial firing rate (D-cells: Saline vs. Morphine, df = 5, F-ratio = 1.034, P = 0.397; H-cells: Saline vs Morphine, df = 11, F-ratio = 1.032, P = 0.419) and final firing rates (D-cells: Saline vs. Morphine, df = 11, F-ratio = 0.365, P = 0.969); H-cells: Saline vs. Morphine, df = 11, F-ratio = 0.333, P = 0.978). Bonferroni-corrected post-hoc tests of firing at each current strength also did not reveal any statistical differences between treatments.

Thus, neither the decrease in soma size not the decrease in excitability of H-type orexin-EGFP neurons following two weeks of daily morphine treatment were still apparent following four weeks of passive withdrawal.

### A single injection of morphine was insufficient to impair excitability of H-type orexin neurons

On the last treatment day, Addiction cohort mice received a final injection of either saline or morphine two hours prior to brain slice preparation. To test for possible lingering effects of morphine from this dose, we injected a different cohort of orexin-EGFP mice (1-day cohort) with a single injection of either saline or morphine (50 mg/Kg) and prepared brain slices two hours later.

We found that there were no significant differences in the passive properties or action potential shape between orexin neurons from saline- and morphine-treated 1-day cohort mice, for either D or H cells (Table 3). However, rheobase was lower by ∼35 % (4 pA), for H-type neurons from morphine-injected mice compared to saline-treated controls (Table 3). Notably, this was opposite to the change in rheobase following two weeks of daily morphine injections, where H-cells required additional depolarizing current to fire a spike.

**Table 3.** Cell type properties from mice after a single saline or morphine treatment. Passive and active properties of D-cells (left columns) and H-cells (right columns) from mice treated with a single injection of saline or morphine (50 mg/kg). Measured parameters are displayed as mean ± SEM. Columns show D-cell data from mice treated with saline (D-type (Saline)) or morphine (D-type (Morphine)). Each measurement is followed by the number of cells (n) measured. The value in the column labelled P is the probability of rejecting the null hypothesis when it is true that the Saline and Morphine values were drawn from the same distribution computed from a two-tailed, two-sample T- test with unequal variance.

|  | Saline D |  | Morphine D |  | P | Saline H |  | Morphine H |  | P |
| --- | --- | --- | --- | --- | --- | --- | --- | --- | --- | --- |
|  | Mean ± SEM | n | Mean ± SEM | n |  | Mean ± SEM | n | Mean ± SEM | n |  |
| Passive Properties |  |  |  |  |  |  |  |  |  |  |
| Capacitance (pF) | 22.52 ± 2.07 | 13 | 22.29 ± 1.42 | 28 | 0.927 | 20.83 ± 1.03 | 16 | 20.77 ± 1.65 | 15 | 0.974 |
| Input Resistance (Gohm) | 1.25 ± 231.68 | 13 | 1.25 ± 1.9 | 28 | 0.992 | 1.2 ± 207.95 | 16 | 1.55 ± 211.45 | 15 | 0.256 |
| Active Properties |  |  |  |  |  |  |  |  |  |  |
| Rheobase (pA) | 13.17 ± 2.51 | 12 | 11.08 ± 1.2 | 26 | 0.464 | 12.38 ± 1.52 | 16 | 8 ± 2.24 | 13 | <0.05 |
| Spontaneous Firing Rate (Hz) | 8.5 ± 1.07 | 12 | 7.96 ± 0.81 | 24 | 0.691 | 10.43 ± 1.74 | 14 | 8.75 ± 1.13 | 14 | 0.428 |
| Threshold (mV) | -40.05 ± 1.39 | 11 | -39.6 ± 0.61 | 26 | 0.773 | -39.74 ± 0.86 | 16 | -38.62 ± 0.76 | 13 | 0.341 |
| Rising Slope Max (mV/ms) | 160.47 ± 9.3 | 11 | 152.84 ± 5.42 | 26 | 0.487 | 186.71 ± 6.55 | 16 | 196.18 ± 12.17 | 13 | 0.502 |
| AP Max (mV) | 52.49 ± 2.05 | 11 | 50.72 ± 1.15 | 26 | 0.461 | 53.42 ± 1.57 | 16 | 54.96 ± 1.17 | 13 | 0.438 |
| AP Amplitude (mV) | 92.54 ± 1.42 | 11 | 90.32 ± 1.19 | 26 | 0.243 | 93.16 ± 1.51 | 16 | 93.59 ± 1.67 | 13 | 0.853 |
| Falling Slope Max (mV/ms) | -72.14 ± 3.62 | 11 | -71.85 ± 2.99 | 26 | 0.951 | -83.43 ± 4.34 | 16 | -94.27 ± 6.14 | 13 | 0.163 |
| Width at Half Max (ms) | 1.49 ± 0.05 | 11 | 1.49 ± 0.04 | 26 | 0.956 | 1.31 ± 0.04 | 16 | 1.22 ± 0.05 | 13 | 0.193 |
| AHP Min (mV) | -67.51 ± 1.16 | 11 | -67.88 ± 0.6 | 26 | 0.781 | -70.04 ± 1.10 | 16 | -70.79 ± 0.91 | 13 | 0.603 |

Most of both D- and H-cells displayed a “continuous” firing pattern following a single saline or morphine injection (Fig 8A) and there was no significant impact of the morphine injection on the distribution of firing patterns expressed by D- or H-cells (D-cells: X^2^ = 3.75, df = 2, N = 38, P = 0.384; H- cells: X^2^ = 4.13 (df = 2, N = 30, P = 0.131). There was also no evidence that a single dose of morphine impaired repetitive firing of H-cells (Fig 8 B right, C right). Rather, we found that the initial, but not final firing rates were modestly increased (10-15 %) at a few current strengths for H- and D- type neurons (Fig 8B, C) as determined by Bonferroni post-hoc testing following 2-way rmANOVAs of D- and H-cell firing rates with treatment (Saline vs. Morphine), current strength and their interaction (treatment * current) as factors. For H-cells, neither the initial firing rate rmANOVA was significant for treatment (df = 1, F-ratio = 2.35, P = 0.139) or interaction (df = 11, F-ratio = 0.614, P = 0.816), nor was the final firing rate significant for either treatment (df = 1, F-ratio = 0.680, P = 0.418) or interaction (df = 11, F-ratio = 0.734, P = 0.706). Nevertheless, post-hoc testing identified the Initial firing rate of H-cells to be higher following a single morphine injection at 40, 50, 70, 90 and 100 pA (Fig 8 B, right). For D-cells, neither the initial firing rate (df = 1, F-ratio = 1.478, P = 0.232; Fig 8 B, left) nor final firing rate (df = 1, F-ratio = 0.936, P = 0.340; Fig 8C, left) rmANOVA was significant for treatment but both were significant for the interaction term (Initial rate: df = 11, F-ratio = 2.863, P = 0.001; Final rate: df = 11, F-ratio = 1.852, P = 0.045). Post-hoc testing revealed that the Initial firing rate was higher following morphine treatment at 110 and 120 pA, but that there were no differences at any current steps for final firing rate.

**Figure 8:**
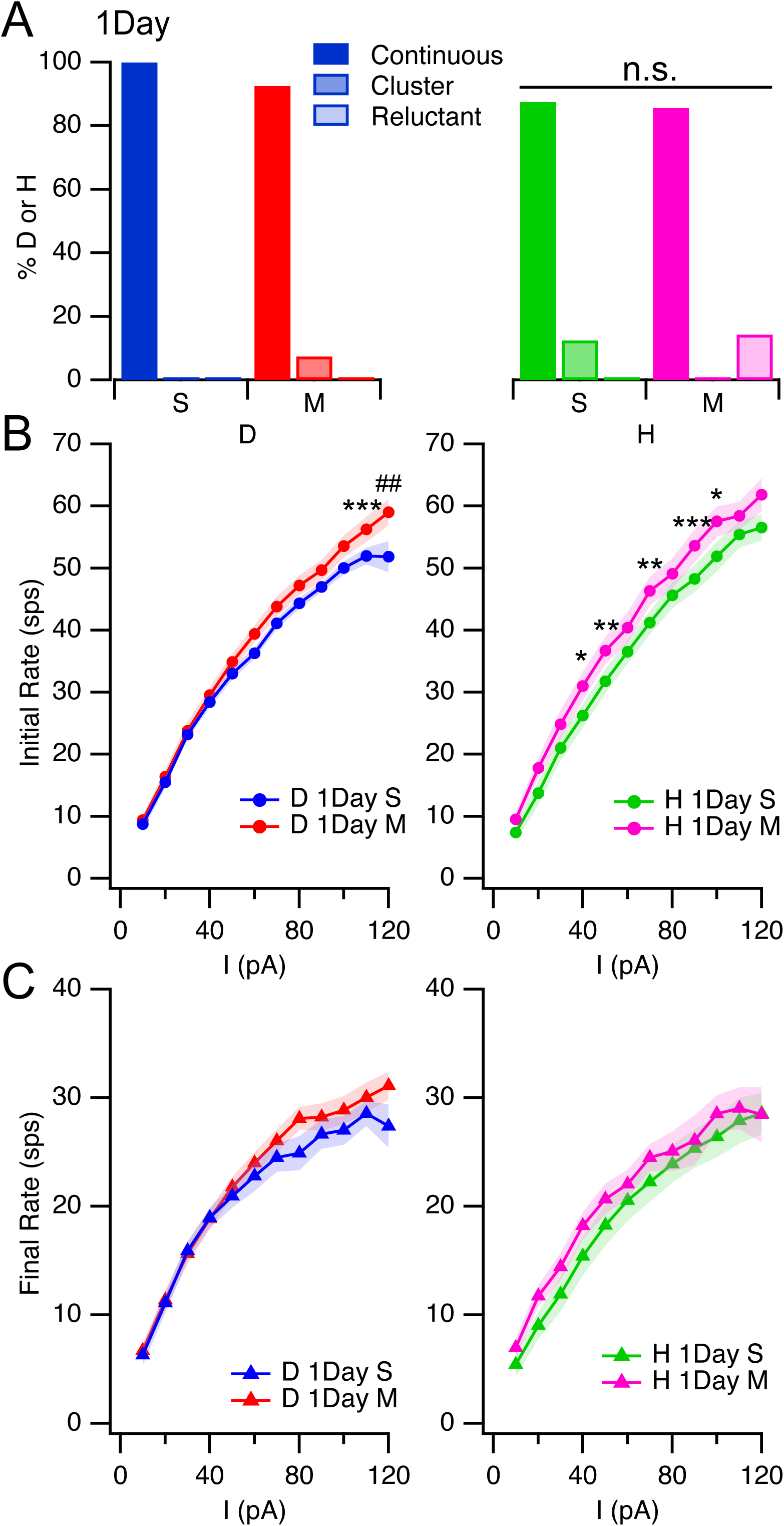
A single morphine injection did not impair firing of D- or H-type orexin neurons. **A.** Distribution of recorded D- (Left) and H-cells (Right) with “continuous”, “cluster” and “reluctant” firing patterns from slices made following a single injection of saline (S, Blue for D-cells; Green for H-cells) or morphine (M, red for D-cells; pink for H-cells). Percent of D and H-type orexin neurons with each repetitive firing pattern. **B, C.** Firing frequency (mean ± SEM) – current (I) curves for initial (B) and final (C) firing rates following a single saline or morphine injection for D-cells (Left) and H-cells (Right). Statistical significance for Bonferroni-corrected pairwise post-hoc tests (* P < 0.05; ** P < 0.01; *** P < 0.005; ## P < 0.0001). Abbreviations: n. s., not significant; S, saline-treated; M, morphine-treated; 1Day, Single injection cohort; sps, spikes/second.

Thus, a single morphine exposure was insufficient to impair spike generation or repetitive firing ability of H-cells. This indicates that the reduced excitability after two weeks of daily morphine injections does not result from the acute effects of morphine lingering from the last injection and that more than a single morphine dose is necessary to induce these changes. Instead, a single injection of morphine appeared to produce a modest increase in excitability of D- and H-cells.

### Chronic morphine exposure downscales the F-I gain of H-type orexin neurons

To compare the within-treatment effects of saline or morphine on the firing ability of D- and H-cells, we superimposed the average F - I curves from the 1-Day, Addiction and Withdrawal cohorts for each treatment. F - I curves were then compared by two-way rmANOVAs with cohort, current and the interaction (cohort *current) as factors. This was followed by Bonferroni post-hoc tests at each current strength. This analysis underscored that while there were only a few pair-wise differences between F - I curves for D-cells from all cohorts (not shown), there were substantial differences between F - I curves for H-cells from cohorts treated with saline and morphine. Fig 9 A illustrates these differences for the final firing rate F - I curves following saline- (left) and morphine- (right) treatment, although initial firing F - I curves showed similar differences (data not shown). The rmANOVA was not different for the saline-treated cohort factor (df = 2, F = 0.89, P = 0.42) or for the interaction term (df = 22, F = 0.88, P = 0.63), but post-hoc testing indicted there was a significant 5 - 6 sps lower average final firing rate for current strengths between 60 - 100 pA for saline-treated H-cells from the Addiction cohort compared to those from the 1D cohort (P < 0.005; Fig 9 A, left). This suggests that daily saline injections also modestly reduced the firing ability of H-cells.

**Figure 9:**
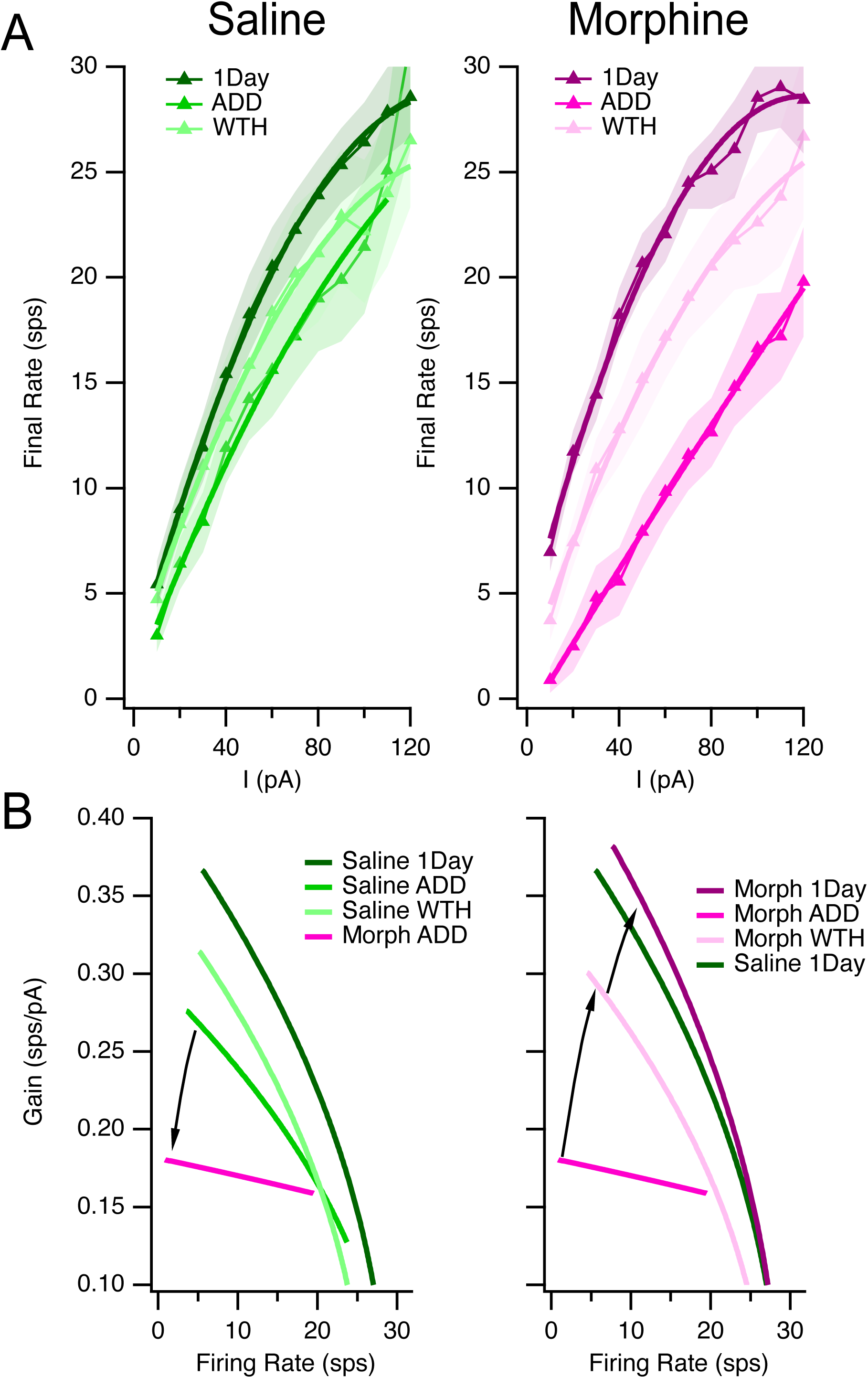
Chronic but not acute morphine exposure downscales firing gain of H-type orexin neurons. **A.** Final firing rate (mean ± SEM) vs. current (I) curves for H-cells are superimposed for saline-treated cohorts of mice (left, green symbols) and morphine-treated mice (right, pink symbols). The 1-day cohort (1Day) is darkest, the Addiction cohort (ADD) is lighter, and the Withdrawal cohort (WTH) is lightest. Corresponding sold lines are best-fit second order polynomials. **B.** Comparison of mean firing rate gain vs. mean firing rate for each cohort. The left graph compares the gain-firing relation for morphine-treated H-cells from the Addiction cohort to those from the saline-treated H-cells from each cohort. The downward arrow connects the gain-firing curve from saline-treated and morphine-treated cells from the Addiction cohort to illustrate the downscaling produced by morphine treatment. Right graph compares the gain-firing relation for morphine-treated H-cells from the Addiction cohort to those from the morphine-treated H-cells from the other cohorts. The long upward arrow connects the curve from the morphine-treated Addiction cohort to that of the Morphine-treated withdrawal cohort to illustrate the recovery from downscaling following 4 weeks of morphine abstinence. The shorter arrow connects the Withdrawal cohort curve to the 1-day morphine cohort curve to illustrate that after 4 weeks of morphine abstinence, the gain-firing relation is still reduced. The curve from the saline-treated mice from the 1-day cohort is shown in each graph for reference.

For the morphine-treated H-cell cohorts (Fig 9 A right), the final firing rate rmANOVA was significant for the cohort factor (df = 2, F = 8.59, P = 0.001), but not for the cohort * current interaction (df = 22, F = 1.25, P = 0.21). However, post-hoc testing between the 1-Day and the Addiction cohorts revealed large reductions in mean firing rate of ∼6 - 13 sps at each current strength (p < 1 x 10^-6^ for each current). Post-hoc testing also found that the mean firing rates of H-cells in the Addiction cohort were 6-8 sps lower than that from the Withdrawal cohort over most current strengths (30 – 110 pA; p < 0.01). There were also smaller (∼ 5-6 sps), but still significant differences between 1-Day and Withdrawal cohorts over a range of currents (40 – 70 pA, and 100 pA, p < 0.05 for each). Thus, the most depressed F-I curves were expressed by H-cells from mice treated with daily morphine injections for two weeks. In contrast, the F-I curves were most robust for cells from mice in the 1-day cohort who received a single morphine injection, while F-I curves achieved intermediate levels following 4 weeks of withdrawal from two-weeks of morphine treatment.

To estimate the how the shape of these F - I curves differed between treatments, we fit these average curves with second order polynomials (Fig 9A; solid lines). We then estimated the average firing rate gain for each condition by taking the derivative of these best-fit polynomials and compared the gain curves by plotting them against their corresponding firing rates (Fig 9 B). In this representation, purely additive or subtractive changes to the F - I curve, like those resulting from shunting inhibition, which translate the F - I curve along the current axis, result in gain-firing rate curves that are superimposed (Chance, Abbott and Reyes, 2002). Rather than producing a subtractive effect, two weeks of morphine treatment resulted in a striking downscaling in firing gain. Compared to H-cells from mice treated with saline for two weeks, the lowest firing rate for neurons from morphine-treated mice (K_0_) was reduced by 76 % and their coefficients for the linear (K_1_) and squared current terms (K_2_) of the best-fit polynomial was reduced by 35 % and 87 %, respectively (Fig 9B, left panel arrow). These reduced coefficients reflect the lower and nearly constant gain - firing curve for morphine-treated neurons. This is a divisive reduction in firing gain of ∼55 % for small inputs but the magnitude of this gain reduction decreases at higher firing rates where it approaches the firing gain of H-cells from saline-treated mice at their highest firing rates. This predicts that H-cells from morphine-treated mice will have a reduced ability to encode input signals into firing rate changes and that the impairment will be worse for smaller inputs. This representation also illustrates the lower gain-firing relation for H-cells from the saline-treated Addiction cohort compared to the saline-treated H-cells from the 1-day cohort. As noted above, this is consistent with the s.q. injections themselves attenuating the firing gain of H-type orexin neurons.

Following four weeks of withdrawal, neurons from morphine treated mice have gain – firing rate relations that correspond well to that of cells recorded from saline treated mice after four weeks of withdrawal. This indicates that the decreased encoding capability of H-type orexin neurons after chronic morphine exposure is mostly reversed following four weeks of morphine abstinence. Nevertheless, peak gains remain lower than that for neurons recorded after a single dose of morphine or saline (Fig 9 B Right, short arrow).

### D & H-type orexin neurons are intermingled throughout the orexin field

It has been proposed that the orexin cell field is organized into a functional topography (Harris and Aston-Jones, 2006) based, in part, on the finding that orexin neurons lateral to the fornix increase c-fos labelling following the development of a conditioned place preference to morphine (Harris et al., 2007). We therefore examined the distribution of our sampled D- and H-cells with respect to the fornix. We plotted the locations of all neurons we identified from the Addiction, Withdrawal, and 1-Day cohorts onto four corresponding atlas pages (Franklin and Paxinos, 2008) from bregma −1.58 mm to −1.94 mm according to their distances from the fornix and mammillothalamic tract (Fig 10A). We then bisected the orexin field into medial and lateral parts with a line through the fornix to determine their distribution on either side. This showed that while ∼60% of our recorded cells were lateral to the fornix, both cell types were distributed throughout the orexin field (Fig 10B). A similar proportion of D- and H-cells were sampled on either side of the fornix, with ∼57 % D- and ∼43% H-cells on both the medial (DMH/PFA) and lateral (LH) sides (Fig 10 C). When we examined the anterior-posterior distribution of D- and H-cells (Fig 10 D), we found that H-cells were enriched lateral to the fornix in the most rostral section while D-cells were enriched lateral to the fornix in the next two more posterior sections. Thus, while D and H-cells are present throughout the orexin field, H cells are overrepresented in the most rostral LH and D-cells are overrepresented more caudally in the LH.

**Figure 10.**
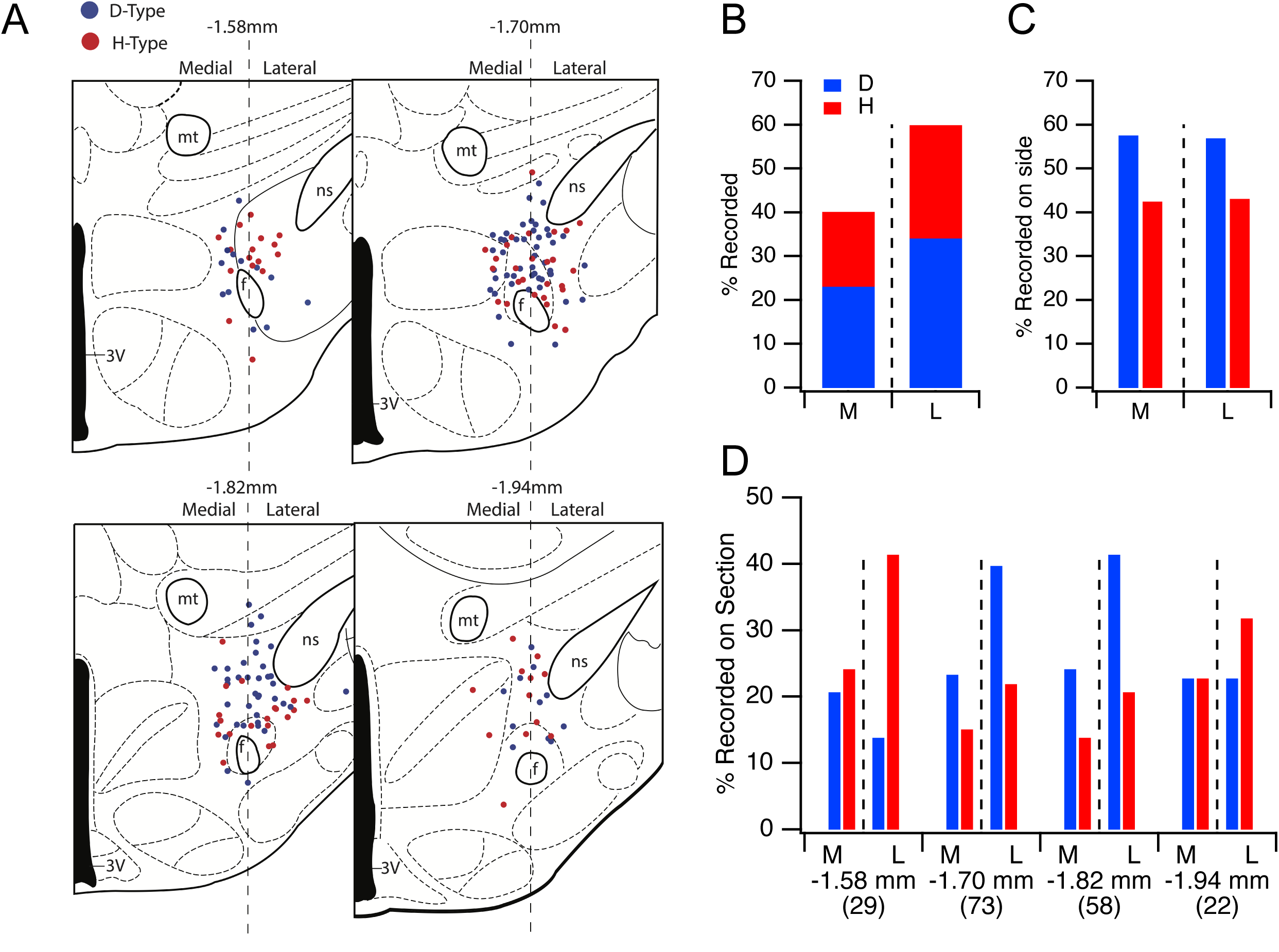
D- and H- type orexin neurons were intermingled throughout the orexin neuron field. **A.** Recorded D- (blue symbols) and H-cells (red symbols) were mapped onto four atlas sections spanning from −1.58 mm to −1.94 mm from bregma. Dotted line bisects the fornix to divide each section into medial and lateral regions. **B.** Percent of recorded cells (n = 182) medial (M) and lateral (L) to the fornix. **C.** Percent of cells recorded on each side that were identified as D- or H-cells. **D.** Anterior to posterior distribution of D- and H- cells medial and lateral to the fornix as a percent of cells in each section. Number of cells mapped to each section indicated in parentheses. Abbreviations: 3V, third ventricle; mt, mammillothalamic tract; f, fornix; ns, nigrostriatal tract.

## Discussion

In this study we found that chronic morphine decreased orexin-EGFP soma size, but did not decrease C_m_ or increase in R_m_. Instead, we found a impairment in excitability of H-type orexin neurons while D-type orexin neurons were entirely spared. A single morphine injection was insufficient to induce this impairment, and four weeks of passive withdrawal was sufficient to reverse the impaired H-cell excitability and soma size decrease. Finally, D- and H-cells were distributed throughout the orexin cell field, but we found H-cells were enriched rostrally, while D-cells were enriched more caudally within the LH.

### Somatic structural plasticity resulting from chronic morphine exposure

Chronic exposure to drugs of abuse produces well-documented structural changes in spines, dendrites and somata of reward circuitry neurons (Robinson and Kolb, 2004; Russo et al., 2010). The effect of chronic morphine on orexin soma size appears similar to its effect on VTA DA neurons. Notably, chronic morphine, but not ethanol or psychostimulants, decreases soma size of VTA dopamine neurons and increases spontaneous firing *in vitro* and *in vivo* (Sklair-Tavron et al., 1996; Mazei-Robison et al., 2014; Simmons, Wheeler and Mazei-Robison, 2019). Moreover, firing rate and soma size are tightly linked in VTA DA neurons, since augmenting firing with a dominant-negative K^+^ channel subunit (KCNAB2) decreased soma size, while suppressing firing with Kir2.1 prevented morphine-induced soma size decreases (Mazei-Robison et al., 2011; Koo et al., 2012). However, unlike VTA DA neurons, morphine directly inhibits ∼50% of orexin neurons *in vitro* (Li and van den Pol, 2008) and chronic morphine did not alter spontaneous firing of orexin-EGFP neurons *in vitro*, suggesting a different mechanism may be involved in orexin neurons. Nevertheless, systemic morphine was reported to induce sustained firing of putative orexin neurons *in vivo* (Thannickal et al., 2018) which may suffice to drive the soma size decrease in orexin neurons if similar signaling pathways are engaged by chronic morphine. Future experiments should determine if orexin neurons are regulated by such activity-dependent processes.

### H- but not D-type orexin neurons showed impaired excitability following chronic morphine

Changes in intrinsic excitability have been described throughout the reward system following both repeated non-contingent and contingent drug exposure (Kourrich, Calu and Bonci, 2015; Hearing, 2019). Multiple mechanisms are involved and vary depending on cell-type (Kourrich and Thomas, 2009; Kim et al., 2011) and conditions (Mu et al., 2010). The impaired excitability we found in H-type orexin neurons included changes in spike generation that were consistent with attenuated voltage-gated sodium currents (increased rheobase, reduced spike amplitude and rise-times, that were exacerbated with sequential spikes). These effects are similar to changes reported in ventral subiculum neurons during amphetamine withdrawal (Cooper et al., 2003) and in medium spiny NAc neurons during cocaine withdrawal (Zhang, Hu and White, 1998). Numerous other currents have also been linked to impaired repetitive firing (irregular and slower firing) following chronic drug exposure (Ishikawa et al., 2009; Chen et al., 2013; Kourrich et al., 2013; Leyrer-Jackson, Hood and Olive, 2021) and are among the numerous candidates underlying impaired H-cell excitability.

### Functional roles and hypothalamic distribution of orexin neurons

Selective excitability impairment of H-cells by chronic morphine may indicate that they are more engaged in reward processing than D-cells. Conversely, D-cells may be more engaged with regulating arousal, since six hour of sleep deprivation selectively increased their excitability (Briggs et al., 2019).

Importantly, both findings support the idea that D- and H-cells are functionally specialized, as originally proposed based on differences in glucose sensing (Williams et al., 2008a). It has been proposed that LH orexin neurons lateral to the fornix regulate reward processing and preferentially project to the VTA and NAc, while PFA/DMH orexin neurons medial to the fornix regulate arousal and stress and preferentially project to LC, TMN and LDT/PPT (Harris and Aston-Jones, 2006). However, D- and H-cells were intermixed throughout the entire orexin field, as previously observed (Williams et al., 2008a). We did find that H-cells were enriched in the most rostral LH region and D-cells were enriched more caudally in the LH. This implies that rostral LH orexin neurons expressing increased c-fos to reward cues (Harris, Wimmer and Aston-Jones, 2005; Fragale et al., 2019; James et al., 2019) would be enriched in H-cells while more caudal LH orexin neuron may be enriched in D-cells. Nevertheless, while orexin neurons projecting to the LC and/or TMN appear distinct from those projecting to the VTA and/or NAc, these neurons are intermingled without medial-lateral segregation (Iyer et al., 2018). Moreover, both D- and H-cells project to VTA and LC, but 80 % of these projections were comprised of D-cells (González et al., 2012), raising the possibility that H-cells preferentially project elsewhere. Identifying those targets would be facilitated by molecular markers, but profiling has yet to distinguish D- or H-cells (Yelin-Bekerman et al., 2015; Mickelsen et al., 2017; Mickelsen et al., 2019; Sagi, de Lecea and Appelbaum, 2021).

### Impaired encoding and decoding by H-cells following chronic morphine

Increased firing rate variability and the downscaling of firing gain diminishes the ability of H-cells to encode synaptic inputs into firing rate changes, with the largest impairment occurring for the smallest inputs (Fig 9). Orexin neurons normally encode both rapid sensory-motor signals (Mileykovskiy, Kiyashchenko and Siegel, 2005; Hassani et al., 2016; Karnani et al., 2020) and slower signals related to arousal, metabolism and reward (Lee, Hassani and Jones, 2005; Mileykovskiy, Kiyashchenko and Siegel, 2005; Williams et al., 2008a; Karnani et al., 2011). Since both initial and steady-state firing gain was reduced, our findings suggest the fidelity of both fast and slower signals would be degraded following chronic morphine.

Impaired firing would also be expected to attenuate the ability of H-cells to release the neuropeptides co-expressed by orexin neurons (for review see Ma et al., 2018). Orexin and dynorphin appear co-packaged into dense-core vesicles (Muschamp et al., 2014) which require higher frequency and longer spike trains for exocytosis than do small vesicles (van den Pol, 2012). Although studies of orexin-mediated synaptic currents are limited (Sears et al., 2013; Schöne et al., 2014), orexin fibers innervating the tuberomammilary histaminergic neurons required prolonged firing at > 10 Hz to generate an orexin-dependent slow EPSP, while AMPA EPSPs followed at all frequencies (Schöne et al., 2014). Following chronic morphine, H-cells required about twice the input current to achieve firing rates (> 10 Hz) consistent with peptide release (Fig 6; ∼30 pA saline-treated vs. ∼60 pA morphine-treated mice). Since orexin and dynorphin have opposing actions whose relative strengths differ by target (Li and van den Pol, 2006; Baimel et al., 2017), impairment of H-cell peptide release could differentially impact their targets if D- and H-cells have divergent connections. This may be especially relevant for connections with VTA DA neurons where optical activation of orexin/dynorphin afferents enhances excitation of a majority of lateral shell-projecting VTA DA neuron but inhibits activity in a majority of medial shell- and amygdala-projecting DA neurons (Mohammadkhani, Qiao and Borgland, 2024). If D- and H-cells differentially innervate these targets, impaired peptide release from H-cells could alter the balance of activity in these DA output circuits which may promote motivation for drug seeking.

Orexin system adaptations following chronic morphine and cocaine, including increased neurons (Thannickal et al., 2018; James et al., 2019) with enhanced innervation of LC (McGregor et al., 2022) and VTA (McGregor et al., 2024), are consistent with greater peptide biosynthesis and release. OX1 signaling sustains motivation for cocaine and fentanyl consumption without altering their reward value (Bernstein et al., 2018; Fragale et al., 2019; James et al., 2019). Intriguingly, enhanced motivation for drug consumption resulting from use of an intermittent access (IA) behavioral economics paradigm, further augmented the numbers of orexin neurons and increased the sensitivity of the drug demand curves to the OX1 antagonist SB-334867, requiring only 30% of the dose needed under lower motivation conditions to attenuate demand (James et al., 2019; Fragale, James and Aston-Jones, 2021). This was interpreted to mean that high motivation drug seeking had become more orexin-dependent, perhaps due to greater abundance and release of orexin peptide (James and Aston-Jones, 2022). However, our findings may suggest an alternative interpretation: Perhaps the augmented immunostaining of orexin somata and terminals, representing the “orexin reserve”, results from peptide accumulation in H-cells from decreased release due to impaired excitability. Decreased release might also explain the greater efficacy of antagonists under conditions of high motivation, since there would be lower orexin receptor occupancy and less competition for antagonist. Future research should address this possibility by measuring peptide release, excitation-release coupling and orexin receptor efficacy.

In summary, our findings demonstrate that daily morphine exposure produces a selective, accumulating, and reversible impairment of intrinsic excitability in H-type orexin neurons while sparing D-type neurons recorded from the same slices. Morphine treatment also decreased orexin soma size, which, like H-cell excitability, recovered over 4 weeks of passive withdrawal. Our findings reinforce the idea that orexin neurons with D- and H- response properties are functionally distinct and suggest that the ability of H-cells to encode inputs and transform them into orexin and dynorphin release becomes impaired with morphine dependence. We speculate that these changes contribute to dysregulating the “orexin reserve” and may promote drug seeking.

## Acknowledgements

Supported by NIH grants R01DA034748 and R01NS027881. We thank Drs, Luis De Lecea and William Joseph Giardino for generously providing Hypocretin/Orexin-EGFP founder mice.

## References

Adamantidis AR, Zhang F, Aravanis AM, Deisseroth K, de Lecea L (2007) Neural substrates of awakening probed with optogenetic control of hypocretin neurons. Nature 450:420–424.

Baimel C, Borgland SL (2015) Orexin Signaling in the VTA Gates Morphine-Induced Synaptic Plasticity. J Neurosci 35:7295–7303.

Baimel C, Lau BK, Qiao M, Borgland SL (2017) Projection-Target-Defined Effects of Orexin and Dynorphin on VTA Dopamine Neurons. Cell Rep 18:1346–1355.

Bernstein DL, Badve PS, Barson JR, Bass CE, Espana RA (2018) Hypocretin receptor 1 knockdown in the ventral tegmental area attenuates mesolimbic dopamine signaling and reduces motivation for cocaine. Addict Biol 23:1032–1045.

Borgland SL, Taha SA, Sarti F, Fields HL, Bonci A (2006) Orexin A in the VTA is critical for the induction of synaptic plasticity and behavioral sensitization to cocaine. Neuron 49:589–601.

Boutrel B, Kenny PJ, Specio SE, Martin-Fardon R, Markou A, Koob GF, de Lecea L (2005) Role for hypocretin in mediating stress-induced reinstatement of cocaine-seeking behavior. Proc Natl Acad Sci U S A 102:19168–19173.

Briggs C, Hirasawa M, Semba K (2018) Sleep Deprivation Distinctly Alters Glutamate Transporter 1 Apposition and Excitatory Transmission to Orexin and MCH Neurons. J Neurosci 38:2505–2518.

Briggs C, Bowes SC, Semba K, Hirasawa M (2019) Sleep deprivation-induced pre- and postsynaptic modulation of orexin neurons. Neuropharmacology 154:50–60.

Brown RM, Khoo SY, Lawrence AJ (2013) Central orexin (hypocretin) 2 receptor antagonism reduces ethanol self-administration, but not cue-conditioned ethanol-seeking, in ethanol-preferring rats. Int J Neuropsychopharmacol 16:2067–2079.

Chance FS, Abbott LF, Reyes AD (2002) Gain modulation from background synaptic input. Neuron 35:773–782.

Chemelli RM, Willie JT, Sinton CM, Elmquist J, Scammell TE, Lee C, Richard JA, Williams C, Xiong Y, Kisanuki Y, Fitch TE, Nakazato M, Hammer RE, Saper CB, Yanagisawa M (1999) Narcolepsy in orexin knockout mice: Molecular genetics of sleep regulation. Cell 98:437–451.

Chen BT, Yau HJ, Hatch C, Kusumoto-Yoshida I, Cho SL, Hopf FW, Bonci A (2013) Rescuing cocaine-induced prefrontal cortex hypoactivity prevents compulsive cocaine seeking. Nature 496:359–362.

Cooper DC, Moore SJ, Staff NP, Spruston N (2003) Psychostimulant-induced plasticity of intrinsic neuronal excitability in ventral subiculum. J Neurosci 23:9937–9946.

Cvetkovic-Lopes V, Eggermann E, Uschakov A, Grivel J, Bayer L, Jones BE, Serafin M, Mühlethaler M (2010) Rat hypocretin/orexin neurons are maintained in a depolarized state by TRPC channels. PLoS One 5:e15673.

de Lecea L, Kilduff TS, Peyron C, Gao X, Foye PE, Danielson PE, Fukuhara C, Battenberg EL, Gautvik VT, Bartlett FS, Frankel WN, van den Pol AN, Bloom FE, Gautvik KM, Sutcliffe JG (1998) The hypocretins: hypothalamus-specific peptides with neuroexcitatory activity. Proc Natl Acad Sci U S A 95:322–327.

Eggermann E, Bayer L, Serafin M, Saint-Mleux B, Bernheim L, Machard D, Jones BE, Muhlethaler M (2003) The wake-promoting hypocretin-orexin neurons are in an intrinsic state of membrane depolarization. J Neurosci 23:1557–1562.

Erisir A, Lau D, Rudy B, Leonard CS (1999) Function of specific K(+) channels in sustained high-frequency firing of fast-spiking neocortical interneurons. J Neurophysiol 82:2476–2489.

Fragale JE, James MH, Aston-Jones G (2021) Intermittent self-administration of fentanyl induces a multifaceted addiction state associated with persistent changes in the orexin system. Addict Biol 26:e12946.

Fragale JE, Pantazis CB, James MH, Aston-Jones G (2019) The role of orexin-1 receptor signaling in demand for the opioid fentanyl. Neuropsychopharmacology 44:1690–1697.

Franklin K, Paxinos G (2008) The Mouse Brain in Stereotaxic Coordinates, 3rd Edition: Academic Press.

Gentet LJ, Stuart GJ, Clements JD (2000) Direct measurement of specific membrane capacitance in neurons. Biophys J 79:314–320.

Gentile TA, Simmons SJ, Barker DJ, Shaw JK, Espana RA, Muschamp JW (2018) Suvorexant, an orexin/hypocretin receptor antagonist, attenuates motivational and hedonic properties of cocaine. Addict Biol 23:247–255.

Georgescu D, Zachariou V, Barrot M, Mieda M, Willie JT, Eisch AJ, Yanagisawa M, Nestler EJ, DiLeone RJ (2003) Involvement of the lateral hypothalamic peptide orexin in morphine dependence and withdrawal. J Neurosci 23:3106–3111.

González JA, Jensen LT, Fugger L, Burdakov D (2012) Convergent inputs from electrically and topographically distinct orexin cells to locus coeruleus and ventral tegmental area. Eur J Neurosci 35:1426–1432.

Harris GC, Aston-Jones G (2006) Arousal and reward: a dichotomy in orexin function. Trends Neurosci 29:571–577.

Harris GC, Wimmer M, Aston-Jones G (2005) A role for lateral hypothalamic orexin neurons in reward seeking. Nature 437:556–559.

Harris GC, Wimmer M, Randall-Thompson JF, Aston-Jones G (2007) Lateral hypothalamic orexin neurons are critically involved in learning to associate an environment with morphine reward. Behav Brain Res 183:43–51.

Hassani OK, Krause MR, Mainville L, Cordova CA, Jones BE (2016) Orexin Neurons Respond Differentially to Auditory Cues Associated with Appetitive versus Aversive Outcomes. J Neurosci 36:1747–1757.

Hearing M (2019) Prefrontal-accumbens opioid plasticity: Implications for relapse and dependence. Pharmacol Res 139:158–165.

Horvath TL, Gao XB (2005) Input organization and plasticity of hypocretin neurons: possible clues to obesity’s association with insomnia. Cell Metab 1:279–286.

Hutcheson DM, Quarta D, Halbout B, Rigal A, Valerio E, Heidbreder C (2011) Orexin-1 receptor antagonist SB-334867 reduces the acquisition and expression of cocaine-conditioned reinforcement and the expression of amphetamine-conditioned reward. Behav Pharmacol 22:173–181.

Ishikawa M, Mu P, Moyer JT, Wolf JA, Quock RM, Davies NM, Hu XT, Schluter OM, Dong Y (2009) Homeostatic synapse-driven membrane plasticity in nucleus accumbens neurons. J Neurosci 29:5820–5831.

Iyer M, Essner RA, Klingenberg B, Carter ME (2018) Identification of discrete, intermingled hypocretin neuronal populations. J Comp Neurol 526:2937–2954.

James MH, Aston-Jones G (2020) Introduction to the Special Issue: “Making orexin-based therapies for addiction a reality: What are the steps from here?”. Brain Res 1731:146665.

James MH, Aston-Jones G (2022) Orexin Reserve: A Mechanistic Framework for the Role of Orexins (Hypocretins) in Addiction. Biol Psychiatry 92:836–844.

James MH, Stopper CM, Zimmer BA, Koll NE, Bowrey HE, Aston-Jones G (2019) Increased Number and Activity of a Lateral Subpopulation of Hypothalamic Orexin/Hypocretin Neurons Underlies the Expression of an Addicted State in Rats. Biol Psychiatry 85:925–935.

Kalogiannis M, Hsu E, Willie JT, Chemelli RM, Kisanuki Y, Yanagisawa M, Leonard CS (2011) Cholinergic modulation of narcoleptic attacks in double orexin receptor knockout mice. PLOS ONE 6:1–14.

Karnani MM, Apergis-Schoute J, Adamantidis A, Jensen LT, de Lecea L, Fugger L, Burdakov D (2011) Activation of central orexin/hypocretin neurons by dietary amino acids. Neuron 72:616–629.

Karnani MM, Schone C, Bracey EF, Gonzalez JA, Viskaitis P, Li HT, Adamantidis A, Burdakov D (2020) Role of spontaneous and sensory orexin network dynamics in rapid locomotion initiation. Prog Neurobiol 187:101771.

Kim J, Park BH, Lee JH, Park SK, Kim JH (2011) Cell type-specific alterations in the nucleus accumbens by repeated exposures to cocaine. Biol Psychiatry 69:1026–1034.

Koo JW, Mazei-Robison MS, Chaudhury D, Juarez B, LaPlant Q, Ferguson D, Feng J, Sun H, Scobie KN, Damez-Werno D, Crumiller M, Ohnishi YN, Ohnishi YH, Mouzon E, Dietz DM, Lobo MK, Neve RL, Russo SJ, Han MH, Nestler EJ (2012) BDNF is a negative modulator of morphine action. Science 338:124–128.

Kourrich S, Thomas MJ (2009) Similar neurons, opposite adaptations: psychostimulant experience differentially alters firing properties in accumbens core versus shell. J Neurosci 29:12275–12283.

Kourrich S, Calu DJ, Bonci A (2015) Intrinsic plasticity: an emerging player in addiction. Nat Rev Neurosci 16:173–184.

Kourrich S, Hayashi T, Chuang JY, Tsai SY, Su TP, Bonci A (2013) Dynamic interaction between sigma-1 receptor and Kv1.2 shapes neuronal and behavioral responses to cocaine. Cell 152:236–247.

Kukkonen JP, Leonard CS (2014) Orexin/hypocretin receptor signalling cascades. Br J Pharmacol 171:314–331.

Lee MG, Hassani OK, Jones BE (2005) Discharge of identified orexin/hypocretin neurons across the sleep-waking cycle. J Neurosci 25:6716–6720.

Leonard CS, Kukkonen JP (2014) Orexin/hypocretin receptor signalling: A functional perspective. Br J Pharmacol 171:294–313.

Leyrer-Jackson JM, Hood LE, Olive MF (2021) Drugs of Abuse Differentially Alter the Neuronal Excitability of Prefrontal Layer V Pyramidal Cell Subtypes. Front Cell Neurosci 15:703655.

Li SB, Giardino WJ, de Lecea L (2017) Hypocretins and Arousal. Curr Top Behav Neurosci 33:93–104.

Li Y, van den Pol AN (2006) Differential target-dependent actions of coexpressed inhibitory dynorphin and excitatory hypocretin/orexin neuropeptides. J Neurosci 26:13037–13047.

Li Y, van den Pol AN (2008) Mu-opioid receptor-mediated depression of the hypothalamic hypocretin/orexin arousal system. J Neurosci 28:2814–2819.

Li Y, Gao XB, Sakurai T, van den Pol AN (2002) Hypocretin/Orexin excites hypocretin neurons via a local glutamate neuron-A potential mechanism for orchestrating the hypothalamic arousal system. Neuron 36:1169–1181.

Lin L, Faraco J, Li R, Kadotani H, Rogers W, Lin X, Qiu X, de Jong PJ, Nishino S, Mignot E (1999) The sleep disorder canine narcolepsy is caused by a mutation in the hypocretin (orexin) receptor 2 gene Cell 98:365–376.

Ma S, Hangya B, Leonard CS, Wisden W, Gundlach AL (2018) Dual-transmitter systems regulating arousal, attention, learning and memory. Neurosci Biobehav Rev 85:21–33.

Mahler SV, Moorman DE, Smith RJ, James MH, Aston-Jones G (2014) Motivational activation: a unifying hypothesis of orexin/hypocretin function. Nat Neurosci 17:1298–1303.

Matovic S, Ichiyama A, Igarashi H, Salter EW, Sunstrum JK, Wang XF, Henry M, Kuebler ES, Vernoux N, Martinez-Trujillo J, Tremblay ME, Inoue W (2020) Neuronal hypertrophy dampens neuronal intrinsic excitability and stress responsiveness during chronic stress. J Physiol 598:2757–2773.

Mazei-Robison MS, Appasani R, Edwards S, Wee S, Taylor SR, Picciotto MR, Koob GF, Nestler EJ (2014) Self-administration of ethanol, cocaine, or nicotine does not decrease the soma size of ventral tegmental area dopamine neurons. PLoS One 9:e95962.

Mazei-Robison MS, Koo JW, Friedman AK, Lansink CS, Robison AJ, Vinish M, Krishnan V, Kim S, Siuta MA, Galli A, Niswender KD, Appasani R, Horvath MC, Neve RL, Worley PF, Snyder SH, Hurd YL, Cheer JF, Han MH, Russo SJ, Nestler EJ (2011) Role for mTOR signaling and neuronal activity in morphine-induced adaptations in ventral tegmental area dopamine neurons. Neuron 72:977–990.

McGregor R, Wu MF, Barber G, Ramanathan L, Siegel JM (2011) Highly specific role of hypocretin (orexin) neurons: differential activation as a function of diurnal phase, operant reinforcement versus operant avoidance and light level. J Neurosci 31:15455–15467.

McGregor R, Wu MF, Thannickal TC, Li S, Siegel JM (2024) Opioid-induced neuroanatomical, microglialand behavioral changes are blocked bysuvorexant without diminishing opioidanalgesia. Nature Mental Health.

McGregor R, Wu MF, Holmes B, Lam HA, Maidment NT, Gera J, Yamanaka A, Siegel JM (2022) Hypocretin/Orexin Interactions with Norepinephrine Contribute to the Opiate Withdrawal Syndrome. J Neurosci 42:255–263.

Mickelsen LE, Kolling FW, Chimileski BR, Fujita A, Norris C, Chen K, Nelson CE, Jackson AC (2017) Neurochemical Heterogeneity Among Lateral Hypothalamic Hypocretin/Orexin and Melanin-Concentrating Hormone Neurons Identified Through Single-Cell Gene Expression Analysis. eNeuro 4.

Mickelsen LE, Bolisetty M, Chimileski BR, Fujita A, Beltrami EJ, Costanzo JT, Naparstek JR, Robson P, Jackson AC (2019) Single-cell transcriptomic analysis of the lateral hypothalamic area reveals molecularly distinct populations of inhibitory and excitatory neurons. Nat Neurosci 22:642–656.

Mileykovskiy BY, Kiyashchenko LI, Siegel JM (2005) Behavioral correlates of activity in identified hypocretin/orexin neurons. Neuron 46:787–798.

Mohammadkhani A, Qiao M, Borgland SL (2024) Distinct Neuromodulatory Effects of Endogenous Orexin and Dynorphin Corelease on Projection-Defined Ventral Tegmental Dopamine Neurons. J Neurosci 44.

Mu P, Moyer JT, Ishikawa M, Zhang Y, Panksepp J, Sorg BA, Schluter OM, Dong Y (2010) Exposure to cocaine dynamically regulates the intrinsic membrane excitability of nucleus accumbens neurons. J Neurosci 30:3689–3699.

Muraki Y, Yamanaka A, Tsujino N, Kilduff TS, Goto K, Sakurai T (2004) Serotonergic regulation of the orexin/hypocretin neurons through the 5-HT1A receptor. J Neurosci 24:7159–7166.

Muschamp JW, Hollander JA, Thompson JL, Voren G, Hassinger LC, Onvani S, Kamenecka TM, Borgland SL, Kenny PJ, Carlezon WA (2014) Hypocretin (orexin) facilitates reward by attenuating the antireward effects of its cotransmitter dynorphin in ventral tegmental area. Proc Natl Acad Sci U S A 111:E1648–1655.

Peyron C, Tighe D, Van den Pol AN, De Lecea L, Heller C, Sutcliffe JG, Kilduff TS (1998) Neurons containing hypocretin orexin project to multiple neuronal systems. The Journal of Neuroscience 18:9996–10015.

Peyron C et al. (2000) A mutation in a case of early onset narcolepsy and a generalized absence of hypocretin peptides in human narcoleptic brains. Nat Med 6:991–997.

Rao Y, Liu ZW, Borok E, Rabenstein RL, Shanabrough M, Lu M, Picciotto MR, Horvath TL, Gao XB (2007) Prolonged wakefulness induces experience-dependent synaptic plasticity in mouse hypocretin/orexin neurons. J Clin Invest 117:4022–4033.

Rao Y, Mineur YS, Gan G, Wang AH, Liu ZW, Wu X, Suyama S, de Lecea L, Horvath TL, Picciotto MR, Gao XB (2013) Repeated in vivo exposure of cocaine induces long-lasting synaptic plasticity in hypocretin/orexin-producing neurons in the lateral hypothalamus in mice. J Physiol 591:1951–1966.

Robinson TE, Kolb B (2004) Structural plasticity associated with exposure to drugs of abuse. Neuropharmacology 47 Suppl 1:33–46.

Russo SJ, Dietz DM, Dumitriu D, Morrison JH, Malenka RC, Nestler EJ (2010) The addicted synapse: mechanisms of synaptic and structural plasticity in nucleus accumbens. Trends Neurosci 33:267–276.

Sagi D, de Lecea L, Appelbaum L (2021) Heterogeneity of Hypocretin/Orexin Neurons. Front Neurol Neurosci 45:61–74.

Sakurai T (2007) [Regulatory mechanism of sleep/wakefulness states by orexin]. Tanpakushitsu Kakusan Koso 52:1840–1848.

Sakurai T, Amemiya A, Ishii M, Matsuzaki I, Chemelli RM, Tanaka H, Williams C, Richard JA, Kozlowski G, Wilson S, Arch J, Bergsma DJ, Yanagisawa M (1998) Orexins and Orexin Receptors- a family of hypothalamic neuropeptides and g protein coupled receptors that regulate feeding behav. Cell Vol 92:573–585.

Scammell TE (2003) The neurobiology, Diagnosis, and Treatment of Narcolepsy. Ann Neurol 53:154–166.

Schindelin J, Arganda-Carreras I, Frise E, Kaynig V, Longair M, Pietzsch T, Preibisch S, Rueden C, Saalfeld S, Schmid B, Tinevez JY, White DJ, Hartenstein V, Eliceiri K, Tomancak P, Cardona A (2012) Fiji: an open-source platform for biological-image analysis. Nat Methods 9:676–682.

Schöne C, Venner A, Knowles D, Karnani MM, Burdakov D (2011) Dichotomous cellular properties of mouse orexin/hypocretin neurons. J Physiol 589:2767–2779.

Schöne C, Apergis-Schoute J, Sakurai T, Adamantidis A, Burdakov D (2014) Coreleased orexin and glutamate evoke nonredundant spike outputs and computations in histamine neurons. Cell Rep 7:697–704.

Sears RM, Fink AE, Wigestrand MB, Farb CR, de Lecea L, Ledoux JE (2013) Orexin/hypocretin system modulates amygdala-dependent threat learning through the locus coeruleus. Proc Natl Acad Sci U S A 110:20260–20265.

Seifinejad A, Ramosaj M, Shan L, Li S, Possovre ML, Pfister C, Fronczek R, Garrett-Sinha LA, Frieser D, Honda M, Arribat Y, Grepper D, Amati F, Picot M, Agnoletto A, Iseli C, Chartrel N, Liblau R, Lammers GJ, Vassalli A, Tafti M (2023) Epigenetic silencing of selected hypothalamic neuropeptides in narcolepsy with cataplexy. Proc Natl Acad Sci U S A 120:e2220911120.

Shaw JK, Ferris MJ, Locke JL, Brodnik ZD, Jones SR, Espana RA (2017) Hypocretin/orexin knock-out mice display disrupted behavioral and dopamine responses to cocaine. Addict Biol 22:1695–1705.

Simmons SC, Wheeler K, Mazei-Robison MS (2019) Determination of circuit-specific morphological adaptations in ventral tegmental area dopamine neurons by chronic morphine. Mol Brain 12:10.

Sklair-Tavron L, Shi WX, Lane SB, Harris HW, Bunney BS, Nestler EJ (1996) Chronic morphine induces visible changes in the morphology of mesolimbic dopamine neurons. Proc Natl Acad Sci U S A 93:11202–11207.

Stuber GD, Wise RA (2016) Lateral hypothalamic circuits for feeding and reward. Nat Neurosci 19:198–205.

Tan Y, Hang F, Liu ZW, Stoiljkovic M, Wu M, Tu Y, Han W, Lee AM, Kelley C, Hajos M, Lu L, de Lecea L, De Araujo I, Picciotto MR, Horvath TL, Gao XB (2020) Impaired hypocretin/orexin system alters responses to salient stimuli in obese male mice. J Clin Invest 130:4985–4998.

Taylor AL (2012) What we talk about when we talk about capacitance measured with the voltage-clamp step method. J Comput Neurosci 32:167–175.

Thannickal TC, Moore RY, Nienhuis R, Ramanathan L, Gulyani S, Aldrich M, Cornford M, Siegel JM (2000) Reduced number of hypocretin neurons in human narcolepsy. Neuron 27:469–474.

Thannickal TC, John J, Shan L, Swaab DF, Wu MF, Ramanathan L, McGregor R, Chew KT, Cornford M, Yamanaka A, Inutsuka A, Fronczek R, Lammers GJ, Worley PF, Siegel JM (2018) Opiates increase the number of hypocretin-producing cells in human and mouse brain and reverse cataplexy in a mouse model of narcolepsy. Sci Transl Med 10.

Ting JT, Daigle TL, Chen Q, Feng G (2014) Acute brain slice methods for adult and aging animals: application of targeted patch clamp analysis and optogenetics. Methods Mol Biol 1183:221–242.

van den Pol AN (2012) Neuropeptide transmission in brain circuits. Neuron 76:98–115.

Williams, Alexopoulos H, Jensen LT, Fugger L, Burdakov D (2008a) Adaptive sugar sensors in hypothalamic feeding circuits. Proc Natl Acad Sci U S A 105:11975–11980.

Williams RH, Alexopoulos H, Jensen LT, Fugger L, Burdakov D (2008b) Adaptive sugar sensors in hypothalamic feeding circuits. Proc Natl Acad Sci U S A 105:11975–11980.

Willie JT, Chemelli RM, Sinton CM, Tokita S, Williams SC, Kisanuki YY, Marcus JN, Lee C, Elmquist JK, Kohlmeier KA, Leonard CS, Richardson JA, Hammer RE, Yanagisawa M (2003) Distinct Narcolepsy Syndromes in Orexn Receptor-2 and Orexin Null Mice: Molecular Genetic Dissection of Non-REM and REM Sleep Regulatory Processes. Neuron 38:715–730.

Yamanaka A, Beuckmann CT, Willie JT, Hara J, Tsujino N, Mieda M, Tominaga M, Yagami K, Sugiyama F, Goto K, Yanagisawa M, Sakurai T (2003) Hypothalamic orexin neurons regulate arousal according to energy balance in mice. Neuron 38:701–713.

Yelin-Bekerman L, Elbaz I, Diber A, Dahary D, Gibbs-Bar L, Alon S, Lerer-Goldshtein T, Appelbaum L (2015) Hypocretin neuron-specific transcriptome profiling identifies the sleep modulator Kcnh4a. Elife 4:e08638.

Zhang XF, Hu XT, White FJ (1998) Whole-cell plasticity in cocaine withdrawal: reduced sodium currents in nucleus accumbens neurons. J Neurosci 18:488–498.

